# Bi-paratopic and multivalent human VH domains neutralize SARS-CoV-2 by targeting distinct epitopes within the ACE2 binding interface of Spike

**DOI:** 10.1101/2020.08.08.242511

**Authors:** Colton J. Bracken, Shion A. Lim, Paige Solomon, Nicholas J. Rettko, Duy P. Nguyen, Beth Shoshana Zha, Kaitlin Schaefer, James R. Byrnes, Jie Zhou, Irene Lui, Jia Liu, Katarina Pance, QCRG Structural Biology Consortium, Xin X. Zhou, Kevin K. Leung, James A. Wells

## Abstract

Neutralizing agents against SARS-CoV-2 are urgently needed for treatment and prophylaxis of COVID-19. Here, we present a strategy to rapidly identify and assemble synthetic human variable heavy (VH) domain binders with high affinity toward neutralizing epitopes without the need for high-resolution structural information. We constructed a VH-phage library and targeted a known neutralizing site, the angiotensin-converting enzyme 2 (ACE2) binding interface of the trimeric SARS-CoV-2 Spike receptor-binding domain (Spike-RBD). Using a masked selection approach, we identified 85 unique VH binders to two non-overlapping epitopes within the ACE2 binding site on Spike-RBD. This enabled us to systematically link these VH domains into multivalent and bi-paratopic formats. These multivalent and bi-paratopic VH constructs showed a marked increase in affinity to Spike (up to 600-fold) and neutralization potency (up to 1400-fold) on pseudotyped SARS-CoV-2 virus when compared to the standalone VH domains. The most potent binder, a trivalent VH, neutralized authentic SARS-CoV-2 with half-minimal inhibitory concentration (IC_50_) of 4.0 nM (180 ng/mL). A cryo-EM structure of the trivalent VH bound to Spike shows each VH domain bound an RBD at the ACE2 binding site, explaining its increased neutralization potency and confirming our original design strategy. Our results demonstrate that targeted selection and engineering campaigns using a VH-phage library can enable rapid assembly of highly avid and potent molecules towards therapeutically important protein interfaces.

## Introduction

The emergence of SARS-CoV-2 and the associated COVID-19 disease has emphasized the need to rapidly generate therapeutics to combat pandemics. SARS-CoV-2 enters cells using the trimeric Spike protein through the interaction of the Spike receptor-binding domain (Spike-RBD) and host angiotensin-converting enzyme-2 (ACE2) on the surface of lung epithelial cells.^1^ Antibody and antibody-like biologics that can block this process are promising therapeutic candidates because of their high specificity and potential neutralization potency.^2^ The majority of antibodies isolated so far against SARS-CoV-2, SARS-CoV-1, and MERS are derived from screening the B-cells of infected patients after viral spread or repurposed from animal immunizations.^3–7^ These approaches, though effective, can be time-consuming and may not necessarily yield neutralizing antibodies. Given the pressing nature of this pandemic, there is a need for multiple additional strategies to rapidly produce potent, recombinant, and neutralizing biologics.

*In vitro* display technologies using yeast or phage are well-established approaches for generating high-affinity binding proteins from large naïve libraries.^8^ *In vitro* selection can be done without the need for infected individuals and only requires the recombinant protein target. One of the recently developed modalities are small single domain antibodies derived from variable heavy homodimer (VHH) domains of antibodies from camels or llamas, often referred to as nanobodies, and are usually obtained by immunization and B-cell cloning.^9–12^ Nanobodies have some advantages. Their single-chain and small size (11 to 15 kDa) allows them to bind epitopes or penetrate tissues that may not be accessible to monoclonal antibodies (mAbs) (150 kDa) and these nanobodies can be rapidly produced in *E. coli*, compared to mammalian expression of two-chain mAbs.^13,14^ However, nanobodies derived from animal immunization can also suffer from long-turnaround times. Although this can be overcome by generating synthetic nanobody libraries,^15,16^ nanobody scaffolds that are animal-derived raise significant concerns regarding immunogenicity if intended for therapeutic purposes. More recently, variable heavy (VH) domains derived from human scaffolds have been produced and tested against a number of targets.^17–19^

Thus, we and others have been interested in developing VH binders to SARS-CoV-2 for the present pandemic, and as a test case for future ones.^20–23^ However, one limitation of synthetic single domain binders is that they often lack the strong binding affinity necessary for therapeutic application. Affinity maturation can improve this, although with a cost of extending the development timeline. Instead, generating linked multivalent or multi-paratopic binders with these VH domains could be a more rapid approach to utilize avidity to boost affinity and efficacy.^24^ VH domains are amenable to linking into such homo- and heterobifunctional formats because they do not present a light chain mispairing challenge like Fabs.^11^

Here, we constructed a human VH-phage library derived from the clinically approved trastuzumab scaffold and validated its use on multiple antigens. By utilizing a masked phage selection strategy, we rapidly identified VH domains at two non-overlapping epitopes within the ACE2 binding site of the SARS-CoV-2 Spike-RBD. By linking these VH domains with a strategic linker into bi-paratopic and multivalent binders, we improved Spike-binding affinity from 10s of nM to ~100 pM without any additional high-resolution structural information. These high affinity binders are capable of potently neutralizing pseudotyped and live SARS-CoV-2. Structural analysis by cryo-electron microscopy (cryo-EM) of the most potent trivalent VH bound to Spike shows that each VH domain precisely targets the ACE2 binding interface on all three RBDs of Spike. We believe our VH-phage library and this multivalent and multi-paratopic approach is highly advantageous when targeted to distinct epitopes within an antigen and can be broadly applied to other viral and non-viral targets to leverage avidity for increased potency.

## Results

### Construction and validation of a synthetic human VH-phage library

To enable the generation of single-domain antibodies against targets such as SARS-CoV-2 Spike, we designed a synthetic VH-phage library using the VH domain (4D5) from the highly stable and clinically successful trastuzumab antibody (**Fig. 1A**).^25,26^ The VH scaffold was modified to include five amino acid changes predicted to reduce aggregation (**Table S1**).^27^ To bias toward colloidal stability, aspartate and arginine or glutamate residues were inserted at the beginning and at two or three terminal positions of CDR H1, as these have been previously used to improve aggregation resistance of VH and scFv fragments from the VH3 germline.^19,28^ Diversity was introduced into CDR H1 and CDR H2 using a minimalistic approach where variability was largely restricted to tyrosine and serine residues **(Fig. 1B)**.^29^ We introduced high-diversity mixtures of amino acids into CDR H3 because it is usually critical to antigen recognition (**Fig. 1B)**, and Fab-phage libraries with highly diverse CDR H3 sequences have successfully generated high-affinity antibodies to a variety of target antigens.^30,31^ Furthermore, charged polar residues such as aspartate were introduced at 10% frequency to decrease net surface hydrophobicity to mitigate aggregation and decrease the propensity for non-specific binders in the library.

**Figure 1:**
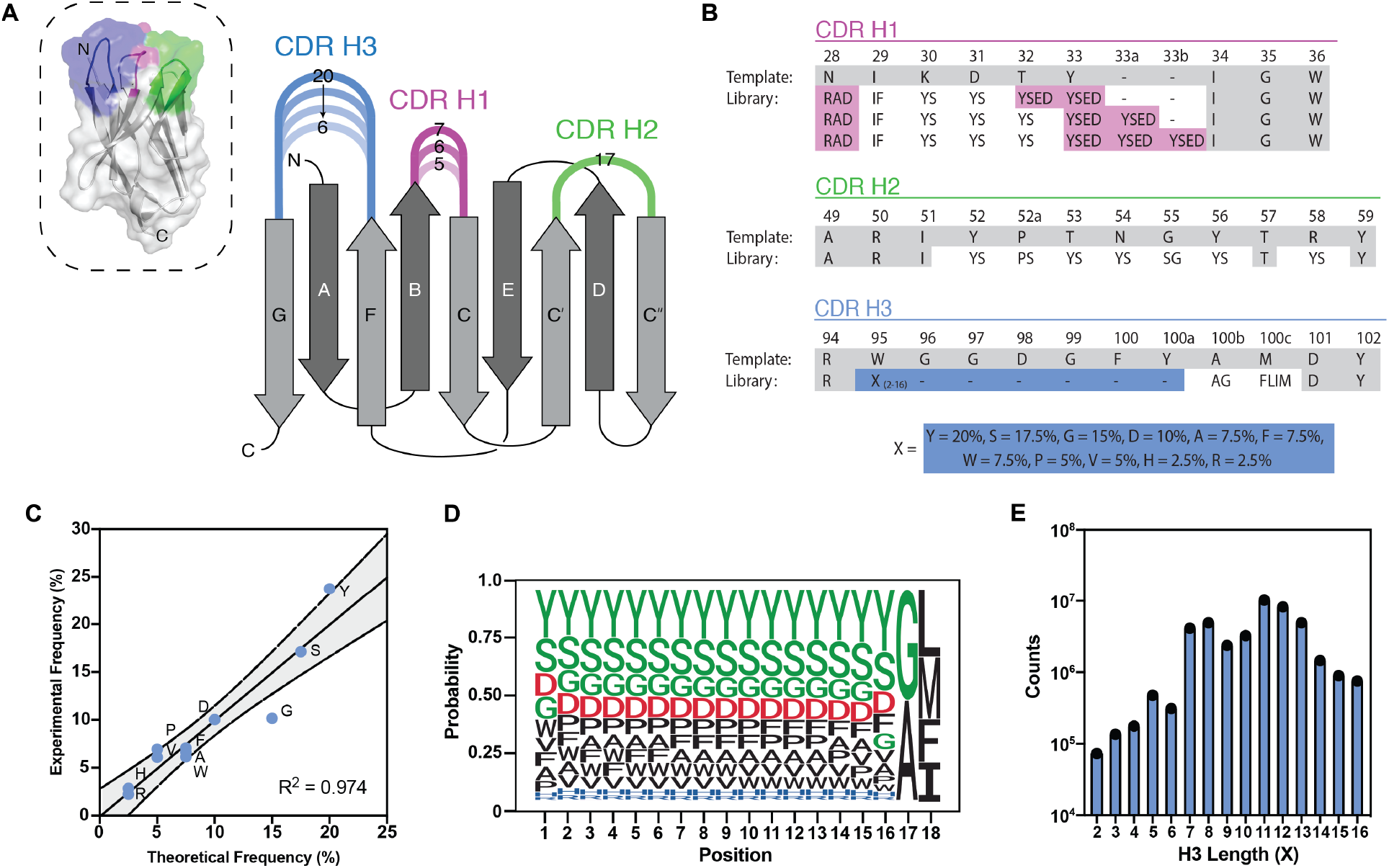
Design and validation of VH-phage library. (**A**) 3D surface representation (left) of the VH-4D5 parental scaffold (PDB:1FVC) and a cartoon diagram (right) where individual CDRs are annotated in color with the designed loop length variations according to Kabat nomenclature.^32^ (**B**) Schematic of CDR amino acid composition as compared to parental template. Positions in pink highlight CDR H1 charged amino acid insertions. Positions in blue highlight insertion of “X” synthetic amino acid mixture. Positions in gray remain unchanged from template. (**C**) NGS analysis of the longest H3 loop (X=18) shows that expected global amino acid frequencies are comparable to designed frequencies. Gray region denotes the 95% confidence interval. (**D**) Representative NGS analysis of the longest H3 loop (X=18) shows positional frequency distribution matches designed frequencies. Position 1 refers to residue 95 (Kabat definition). Data for the other CDR H3 lengths are reported in **Fig. S2**. (**E**) NGS analysis of unique clones shows that all H3 lengths are represented in the pooled VH-phage library.

Based on previous designs, loop length variations of 5 to 7 residues in CDR H1 and 6 to 20 residues in CDR H3 were chosen while CDR H2 was kept constant at 17 residues (Kabat definitions) (**Table S2**).^30,32^ To cover this large sequence space with minimal bias towards different length variants, five separate sub-libraries were constructed by binning CDR H3 loop length insertions (X_2-16_) in incremental sets of three and combined to yield a final library of ~5 x 10^10^ transformants (**Fig. S1**). Analysis of the unique CDR H3 sequences by next-generation sequencing (NGS) show that observed amino acid frequencies closely matched our designs and all CDR H3 length variants were represented in the final library (**Fig. 1C-1E, Fig. S2**). Finally, to test the performance of the library, several rounds of panning were performed on a representative set of six antigens including both cytosolic and membrane proteins. These panning experiments were done in parallel with an in-house Fab-phage library. For all target antigens, high levels of phage enrichment were observed (**Fig. S3**). For the majority of antigens, enrichment levels were comparable or substantially higher for the VH-phage library compared to the Fab-phage library.

### Identification of VH domains that target multiple epitopes within the ACE2 binding site on Spike

To date, most neutralizing mAbs against SARS-CoV-2 target Spike, and not surprisingly many of the most potent target the ACE2 binding interface.^3,7^ While cryo-EM structures show that the ACE2 binding interface remains largely solvent accessible in both the RBD “up” and “down” conformations,^33^ simultaneous intra-molecular engagement by both binding arms of mAbs may be challenging as they are not arranged with the geometries to engage multiple RBDs on a single Spike trimer. Thus, our goal was to target this highly neutralizing epitope with VHs and subsequently link them together to utilize avidity beyond that of a homo-bivalent mAb.

We first expressed the Spike-RBD (residues 328-533) and the ACE2 peptidase domain (residues 1-640) as biotinylated Fc-fusions for VH-phage selections.^34^ To specifically enrich for VH-phage that bind the ACE2 binding site on Spike-RBD, the library was first cleared with the Spike-RBD-Fc/ACE2-Fc complex to remove phage that bind outside the ACE2 binding interface. This was followed by selection on Spike-RBD-Fc alone to enrich for phage that bind the unmasked ACE2 binding site **(Fig. 2A)**. By rounds 3 and 4, significant enrichments for phage that bind Spike-RBD-Fc but not to Spike-RBD-Fc/ACE2-Fc complex were observed **(Fig. S4A)**. Single clones were isolated and characterized for their ability to bind Spike-RBD-Fc by phage enzyme-linked immunosorbent assays (phage-ELISA) **(Fig. S4B)**. Nearly all VH-phage showed enhanced binding to Spike-RBD-Fc over the Spike-RBD-Fc/ACE2-Fc complex, suggesting they bound the same epitope as ACE2 and could potentially block this interaction (**Fig. S4C**). In total, 85 unique VH-phage sequences were identified, and a subset were characterized as recombinant VH domains. We identified three lead VH candidates that bind Spike-RBD with KD values ranging from 23-113 nM (**Fig. 2B-2D, Table 1**). Epitope binning demonstrated that the three VH domains bind at two non-overlapping epitopes we call Site A and Site B within the larger ACE2 binding site (**Fig. S5**). The VH domain that binds Site A (A01) binds independently from the VHs that bind Site B (B01 and B02) (**Fig. 2E-2H**).

**Figure 2:**
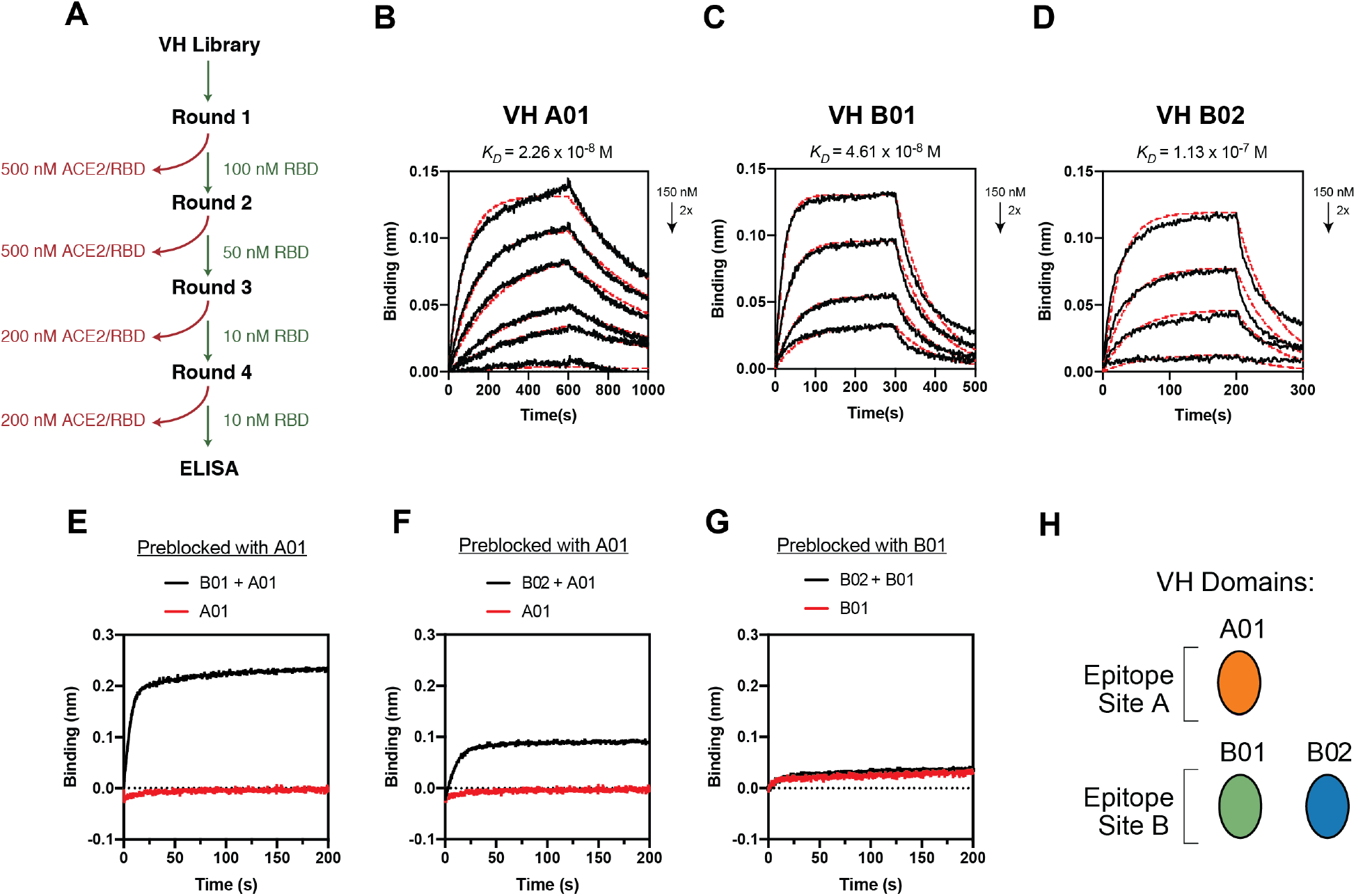
Identification of VH domains that bind Spike-RBD at two unique epitopes by phage display. (**A**) Diagram illustrating phage selection strategy to isolate VH-phage that bind at the ACE2 binding interface. Red indicates clearance of the phage pool by Spike-RBD-Fc/ACE2-Fc complex, green indicates positive selection against Spike-RBD-Fc alone. To increase stringency, successively lower concentrations of Spike-RBD-Fc were used, and after 4 rounds of selection, individual phage clones were analyzed by phage ELISA. BLI of (**B**) VH A01 (**C**) VH B01 and (**D**) VH B02 against Spike-RBD. (**E**) BLI-based epitope binning of VH A01 and VH B01, (**F**) VH A01 and VH B02, (**G**) VH B01 and VH B02. The antigen loaded onto sensor tip was Spike-RBD. (**H**) Diagram of the two different epitope bins targeted by VH domains.

**Table 1:** BLI binding affinity of VH binders to Spike-RBD.

| VH ID | Spike-RBD |  |  |  |
| --- | --- | --- | --- | --- |
| | $K_D$ (nM) | | | |
|  | VH | VH-Fc | VH<sub>2</sub> | VH<sub>3</sub> |
| A01 | 23 | 8.40 | 1.89 | 4.00 |
| B01 | 46 | 0.12 | 0.19 | 0.11 |
| B02 | 113 | 0.19 | 0.46 | 5.18 |
| A01-B01 |  |  | 0.10 |  |
| A01-B02 |  |  | 0.48 |  |
| B01-B02 |  |  | 0.23 |  |

### Bi-paratopic and multivalent VH exhibit dramatically increased affinity to SARS-CoV-2 Spike and block ACE2 binding

We chose two parallel approaches to increase the affinity of the VH binders to Spike through avidity. First, we reasoned that VHs targeting Site A or Site B are in close proximity because they are non-overlapping but compete for the larger ACE2 binding site. Therefore, these VHs could be linked together to engage the same RBD simultaneously and improve affinity through intra-RBD avidity. Using the three VH monomers (A01, B01, B02) as modular units, we generated two bi-paratopic linked dimers (VH_2_) by fusing A01 with B01 or B02 (**Fig. 3A**). In a parallel approach, we aimed to leverage the trimeric nature of Spike and engage multiple RBDs on the same Spike simultaneously to improve affinity through inter-RBD avidity. To that end, we generated mono-paratopic Fc fusions (VH-Fc), linked dimers (VH_2_), and linked trimers (VH_3_) (**Fig. 3A**). The VH_2_ and VH_3_ consisted of a C-to-N terminal fusion of two or three VH monomers via a 20-amino acid Gly-Ser linker (~70 Å) while the VH-Fc consisted of a genetic fusion of VH to the human IgG1 Fc domain via a flexible Fc hinge (~100 Å). The structure of the SARS-CoV-2 Spike trimer suggests the linked VH domains could bridge the distance between RBDs on an individual Spike (<55 Å), but are unlikely to span RBDs between discrete Spike proteins based on the inter-Spike distance on the viral envelope (150-180 Å) (**Fig. S6).**^33,35^

**Figure 3:**
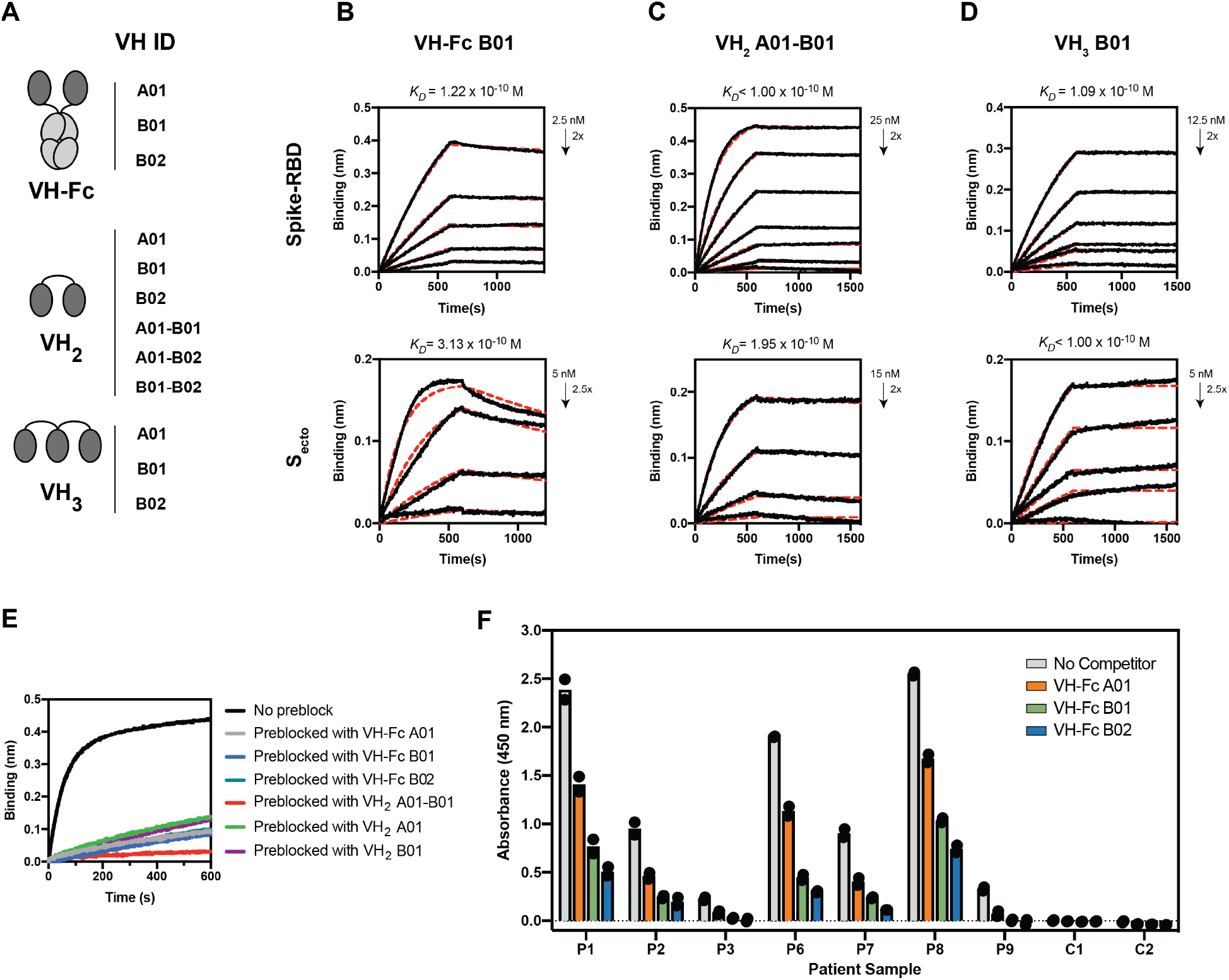
In vitro characterization of multivalent and bi-paratopic VH binders. (**A**) Cartoon depiction of engineered VH binders generated by linking VH domains via Fc-fusion or via a 20-aa Gly-Ser linker. BLI traces of lead VH binders, (**B**) VH-Fc B01, (**C**) VH_2_ A01-B01, (**D**) VH_3_ B01 against RBD (upper panel) or Secto (lower panel). (**E**) Sequential BLI binding experiments that measured binding of ACE2-Fc to S_ecto_ pre-blocked with our VH binders show that multivalent VH binders can block ACE2-Fc binding to _ecto_. (**F**) Competition serology ELISA with convalescent patient sera indicate that VH-Fc binders can compete with patient antibodies. P1-P9 are sera from patients with a history of prior SARS-CoV-2 infection. C1-C2 are two donor sera collected before the SARS-CoV-2 outbreak.

ELISA and BLI binding assays to Spike-RBD show that the VH-Fc, VH_2_, and VH_3_ have 2.7 to 600-fold higher affinity to Spike-RBD (K_D_ = 0.1-8.4 nM) compared to the standalone VH monomers (**Fig. 3B-3D, Table 1, Fig. S7, Fig. S8, Table S3).**Interestingly, fold-increases in affinity were greater for binders that target Site B or both Site A and Site B combined. The most potent constructs bind trimeric Spike ectodomain (S_ecto_) with K_D_s in the hundreds of picomolar range and all utilize VH B01 (**Fig. 3B-3D**). Next, we examined whether these multivalent VH can block ACE2 binding to Spike by testing several high-affinity constructs (VH-Fcs; VH_2_ A01-B01, VH_2_ A01, and VH_2_ B02) in a sequential BLI binding assay. S_ecto_ was immobilized on the biosensor, pre-blocked with each VH binder, and then assayed for binding to ACE2-Fc (**Fig. 3E**). We found that all binders tested substantially blocked binding of ACE2-Fc to S_ecto_. Similarly, we examined whether these engineered VH-Fcs can compete with SARS-CoV-2 Spike-reactive antibodies in convalescent patient serum. Using a competition ELISA format previously developed by our group,^36^ we found that VH-Fcs reduced the binding of patient antibodies to SARS-CoV-2 Spike-RBD (**Fig. 3F**). Taken together these data show that modular reformatting of these VH domains can significantly increase the affinity to the target antigen and block the same immunogenic epitopes as patient-derived Abs.

Lastly, we characterized the biophysical properties of these engineered VH domains by differential scanning fluorimetry (DSF), size exclusion chromatography (SEC), and reconstitution after lyophilization. The VH binders can be rapidly expressed in *E. coli* at high yields (i.e. VH_2_ A01-B01 and VH_3_ B01 express at ~1 g/L in shake flask culture) and have good stabilities (T_m_ = 60-65 °C) (**Fig. S9**). The most potent binders elute as a single mono-disperse peak via SEC (**Fig. S10**), and VH_3_ B01 retains binding to Spike-RBD and a monodisperse SEC profile after lyophilization and reconstitution (**Fig. S11**).

### Bi-paratopic and multivalent VH potently neutralize pseudotyped and live SARS-CoV-2

We then tested the VH binders in pseudotyped virus and authentic SARS-CoV-2 neutralization assays. Pseudotyped virus was used to determine the half-minimal inhibitory concentration (IC_50_) of neutralization for each construct. The VH monomers neutralize pseudotyped virus weakly (IC_50_ > 50 nM), and cocktails of unlinked monomers do not improve potency. In contrast, the multivalent binders (VH_2_, VH_3_, and VH-Fc) neutralize ~10-1000 fold more potently compared to their respective monomeric units (**Fig. 4A, Table 2, Fig. S12**). There was a linear correlation between the in vitro binding affinity (K_D_) to Spike-RBD and the pseudotyped neutralization potency (IC_50_) across the different binders (R^2^ = 0.72) (**Fig. 4B**).

**Figure 4:**
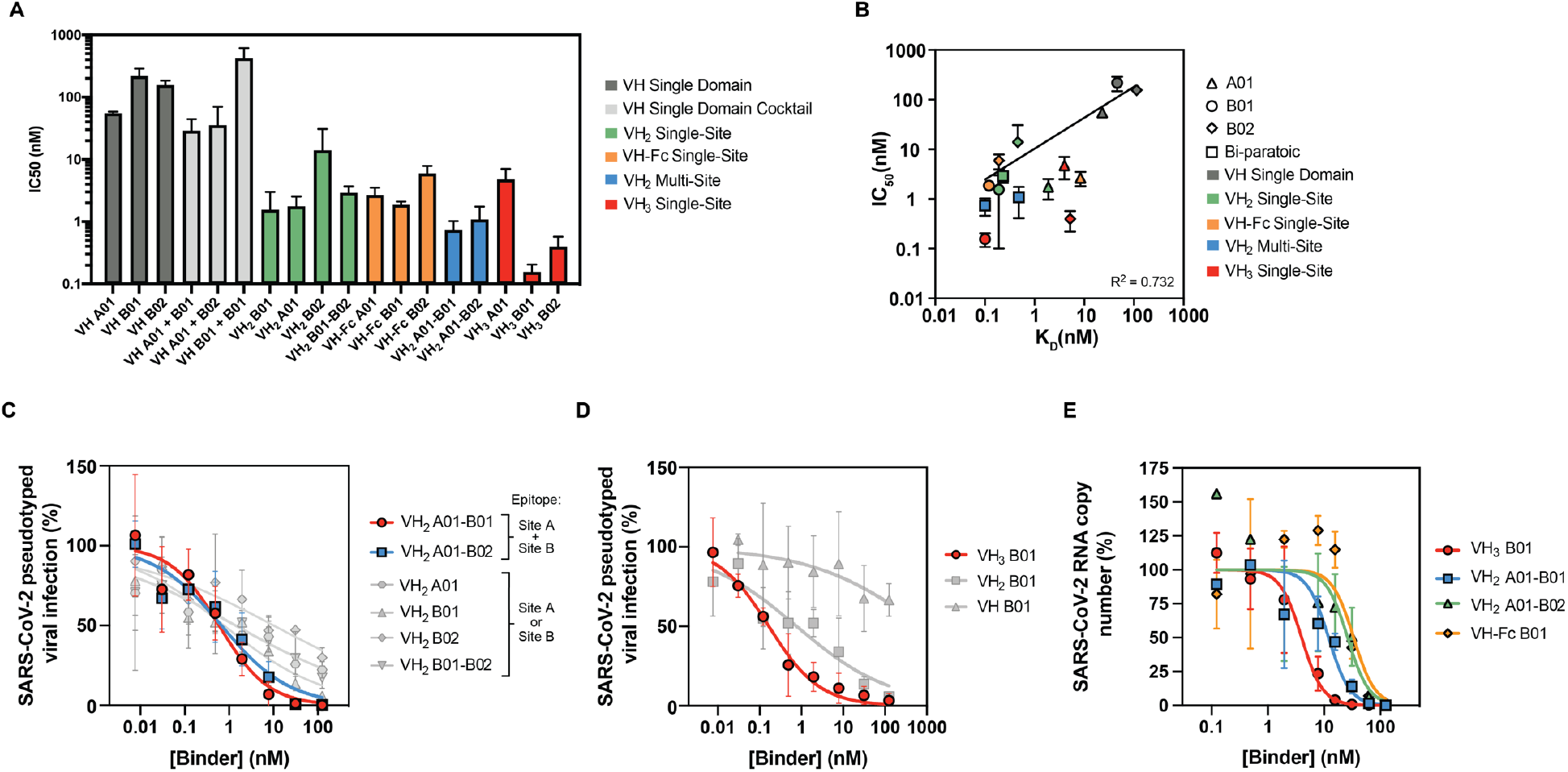
Multivalent and bi-paratopic VH binders neutralize pseudotyped and live SARS-CoV-2. (**A**) Pseudotyped virus IC_50_ of VH binders. Neutralization potency improves when VH domains are engineered into multivalent and bi-paratopic constructs. (**B**) Correlation of in vitro binding affinity (KD) and pseudotyped virus neutralization (IC_50_) of VH binders. Data were fit to a log-log linear extrapolation. (**C**) Pseudotyped virus neutralization curves of multi-site VH_2_ in comparison to single-site VH_2_ demonstrate that the multi-site VH_2_ demonstrate a more cooperative neutralization curve. (**D**) Pseudotyped virus neutralization curves of mono-, bi-, and tri-valent formats of VH B01 demonstrate potency gains driven by valency. (**E**) Authentic SARS-CoV-2 neutralization curves for the most potent VH formats were determined via qPCR of viral genome in cellular RNA. All neutralization data were repeated as two or three biological replicates, with two technical replicates for each condition.

**Table 2:** Pseudotyped virus neutralization IC50.

| VH ID | Pseudotyped virus |  |  |  |
| --- | --- | --- | --- | --- |
|  | IC<sub>50</sub> |  | nM<br>(ng/mL) |  |
|  | VH | VH-Fc | VH<sub>2</sub> | VH<sub>3</sub> |
| A01 | 55.1 ± 3.3<br>(763 ± 46) | 2.67 ± 0.85<br>(209 ± 66) | 1.75 ± 0.77<br>(50.7 ± 22.3) | 4.76 ± 2.28<br>(210 ± 101) |
| B01 | 218 ± 70<br>(3032 ± 978) | 1.86 ± 0.24<br>(145 ± 19) | 1.55 ± 1.45<br>(45.0 ± 42.2) | 0.156 ± 0.048<br>(6.90 ± 2.12) |
| B02 | 156 ± 28<br>(2065 ± 378) | 5.93 ± 1.95<br>(462 ± 152) | 14.0 ± 16.8<br>(388 ± 466) | 0.396 ± 0.176<br>(16.7 ± 7.46) |
| A01 + B01 | 28.9 ± 15.5<br>(400 ± 215) |  |  |  |
| A01 + B02 | 35.3 ± 34.7<br>(491 ± 482) |  |  |  |
| B01 + B01 | 423 ± 193<br>(5608 ± 2554) |  |  |  |
| A01-B01 |  |  | 0.74 ± 0.28<br>(21.4 ± 8.1) |  |
| A01-B02 |  |  | 1.08 ± 0.67<br>(30.6 ± 18.9) |  |
| B01-B02 |  |  | 2.90 ± 0.79<br>(82.4 ± 22.4) |  |

In particular, we observed that bi-paratopic VH_2_ A01-B01 and VH_2_ A01-B02 were stronger neutralizers than unlinked monomer cocktails. Additionally, the neutralization curves of the bi-paratopic (multi-site) VH_2_ differ from the homodimeric (single-site) VH_2_ binding to either Site A or B. That is, the bi-paratopic VH_2_ exhibit a more cooperative transition and fully neutralizes virus, while the homodimeric VH_2_ show a more linear transition and do not fully block viral entry even at high concentrations (**Fig. 4C**). This may reflect mechanistic differences; the bi-paratopic VH_2_ can theoretically engage a single RBD using both VH domains simultaneously (intra-RBD avidity) and more fully occlude the ACE2 binding site, while the homodimeric VH_2_ must bridge separate RBDs within the trimer (inter-RBD avidity).

Furthermore, the increase in neutralization potency as we increase the number of tandem VH units is exemplified by the VH B01-derived binders, where VH_2_ B01 and VH_3_ B01 have IC_50_s that are two to three orders of magnitude lower than the VH B01 monomer (**Fig. 4D**). This is also observed for VH-Fc B01, which also neutralizes two orders of magnitude more potently than the monomer. Interestingly, although the neutralization potency of VH_2_ A01 is better than VH A01, the potency does not improve further when a third domain is added (VH_3_ A01), indicating that epitope-specific geometries can affect the extent to which increasing valency improves potency. The pseudotyped virus neutralization assays demonstrate that the top predicted binders from *in vitro* affinity data, VH_2_ A01-B01, VH_2_ A01-B02, VH_3_ B01, and VH-Fc B01 are indeed the most potent with IC_50_s of 0.74 nM, 1.08 nM, 0.156 nM, and 1.86 nM, respectively.

Lastly, we tested the ability of the most potent VH binders to neutralize authentic SARS-CoV-2 virus. As predicted, VH_3_-B01 was the strongest neutralizer. The VH-Fc B01, VH_2_ A01-B01, and VH_2_ A01-B02 followed trends consistent with both the *in vitro* binding K_D_ and pseudotyped virus IC_50_. VH_3_ B01, VH-Fc B01, VH_2_ A01-B01, and VH_2_ A01-B02 blocked authentic SARS-CoV-2 viral entry with IC_50_s of 3.98 nM, 33.5 nM, 12.0 nM, and 26.2 nM, respectively (**Fig. 4E, Table 3**).

**Table 3:** Live SARS-CoV-2 Neutralization IC50.

| VH ID | SARS-CoV-2 |
| --- | --- |
|  | IC50 nM<br>(ng/mL) |
| VH<sub>2</sub>A01-B01 | 12.0 ± 3.0<br>(347 ± 87) |
| VH<sub>2</sub>A01-B02 | 26.2 ± 6.5<br>(743 ± 184) |
| VH-Fc B01 | 33.5 ± 8.3<br>(2613 ± 647) |
| VH<sub>3</sub>B01 | 3.98 ± 1.56<br>(176 ± 69) |

### Cryo-EM structure of VH_3_ B01 reveals multivalent binding mode

To confirm whether our linking strategy could successfully engage multiple RBDs on Spike, we obtained a 3.2 Å global resolution cryo-EM 3D reconstruction of SARS-CoV-2 S_ecto_ in complex with VH_3_ B01, the most potent neutralizer (**Fig. 5A, Fig. S13-S14**). Although the S2 region of S_ecto_ was resolved at the reported resolution, the RBDs with the bound VH domains were resolved at about 6 Å resolution. However, even at this resolution the structure unambiguously revealed the three RBDs on Spike are in a two “up” and one “down” conformation. Densities corresponding to each VH domain are present on all three RBDs, indicating that VH B01 can bind both “up” and “down” conformations of RBD. SARS-CoV-2 Spike is rarely observed in this conformation with most structures being in all RBD “down” state or one RBD “up” state.^4,6,33,37^ The binding epitope of VH B01 overlaps with the known ACE2 binding site (**Fig. 5B**), confirming the intended mechanism of neutralization and validating the ability of the masked selection strategy to precisely direct a binder toward the intended surface on a target protein.

**Figure 5:**
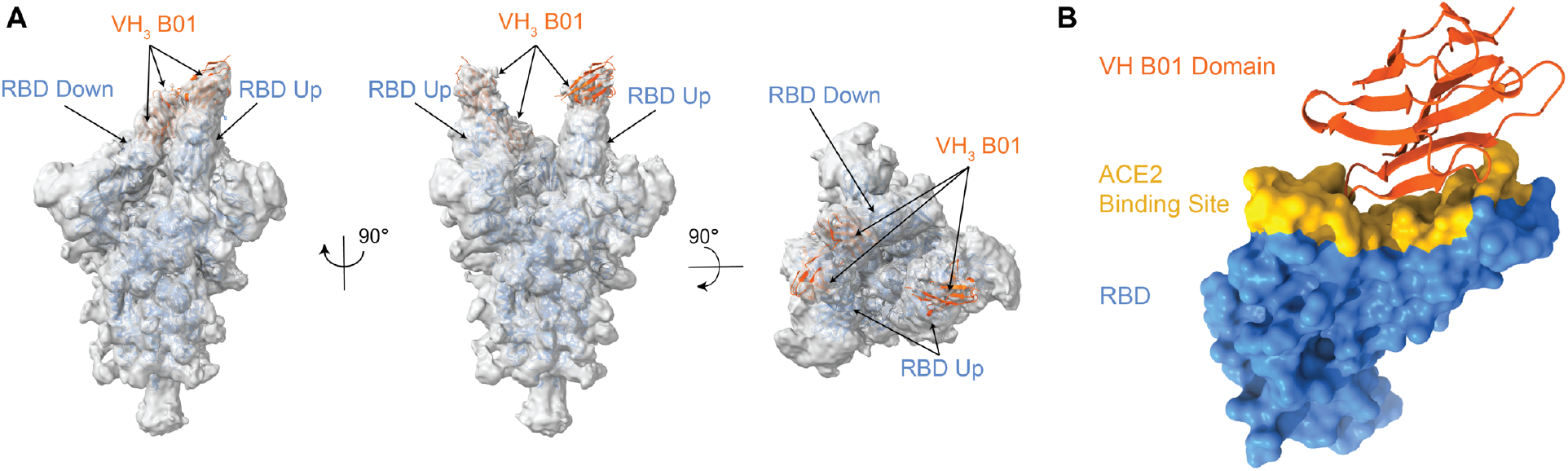
Cryo-EM reveals trivalent VH binding at the ACE2 binding interface of RBD. (**A**) Side and top views of cryo-EM 3D reconstructions of VH_3_ B01 + S_ecto_ are shown with individual VH domain densities of VH_3_ B01 fit with PDB: 3P9W (VH scaffold; orange cartoon). A total of three VH domains, each bound to an RBD of the Spike trimer, are resolved. 3D model of S_ecto_ was fit with reference structure (PDB:6X2B with additional rigid body fit of the individual RBDs; blue cartoon) and shows RBDs in a distinct two “up”, one “down” conformation. Cryo-EM map was low-pass filtered to 6 Å. (**B**) View of the epitope (Site B) of one VH domain from VH_3_ B01. The Site B overlays directly with the ACE2 binding site (yellow surface; contacts defined as RBD residues within 8 Å of an ACE2 residue from PDB:6M0J).

## Discussion

Here we describe a straightforward strategy to rapidly generate linked single-domain binders that potently neutralize SARS-CoV-2. We began by creating and validating a diverse human VH-phage library and generating VH binders to the ACE2 binding interface of Spike-RBD by a masked selection approach. From a panel of 85 unique VH binders, three were identified that bind two separate epitopes within the ACE2 binding interface with nM affinity. In order to bind multiple epitopes simultaneously within an RBD or across RBDs on the Spike trimer, these VH monomers were linked into multivalent and bi-paratopic formats by Gly-Ser linkers or Fc domains without any further high-resolution structural information. This linkage approach not only greatly enhanced affinity, but also substantially improved the viral neutralization efficiency. We confirmed the basis of this potency increase by cryo-EM of the most potent trivalent VH, which blocks the ACE2 binding site on all three RBDs of Spike.

Interestingly, we found there was a difference in the cooperativity of the pseudotyped virus neutralization curves between bivalent VH_2_ that target both Site A and Site B (multi-site) versus VH_2_ that target only Site A or Site B (single-site). Despite similar IC_50_s, the IC95s for these multi-site binders (VH_2_ A01-B01, VH_2_ A01-B02) are much lower than single-site VH_2_ binders. This could indicate a mechanistic difference driven by intra-RBD avidity, as the multi-site VH_2_ can utilize up to 6 binding epitopes on the trimeric Spike, while the single-site VH_2_ are limited to only 3 binding epitopes. Additionally, a more complete occlusion of the ACE2 binding interface (~864 Å^2^)^38^ on Spike-RBD by the multi-site VH_2_ may underlie this difference in neutralization profiles. Although we do not have a structure of VH_2_ A01-B01, we know both A01 and B01 can bind simultaneously within the ACE2 binding site. The cryo-EM structure for B01 shows good coverage of the ACE2 binding site (**Fig. 5B**), while leaving open adjacent space for VH A01 to occupy a non-overlapping epitope.

Additionally, we show that *in vitro* binding affinities and neutralization potencies against this oligomeric target can be dramatically increased with valency. This is exemplified by VH B01, which shows a 460-fold increase in binding affinity and 1400-fold increase in pseudotyped virus neutralization from VH B01 to VH_2_ B01 to VH_3_ B01. A similar trend is observed for VH B02, which also targets the same epitope (Site B). However, VH A01, which targets Site A, is mechanistically distinct as there is no change in potency between a bivalent (VH_2_) and a trivalent (VH_3_) format. This suggests that unlike Site B binders, the trivalent Site A binder may not be able to fully engage all 3 RBDs. This could be due to the specific binding mechanism and epitope of VH A01, conformational differences of the RBDs within the Spike trimer, or spatial constraints of the linker. Structure determination of Site A binders in complex with Spike can elucidate the mechanistic and geometric difference between Site A and Site B epitopes. Additionally, structure-guided approaches to optimize linker lengths and orientations, coupled with an affinity maturation campaign may enable further increases in potency beyond what is demonstrated in this study.

Designed ankyrin repeat proteins (DARPins), llama derived nanobodies, computationally designed proteins, and bivalent Fabs inspired our engineering strategy.^39–45^ These human derived VH domains offers the advantage that they would not require the time-consuming structure-guided humanization process that could be necessary for therapeutic nanobody development. The VH domains also have favorable biophysical properties and scalable production in bacterial systems that is amenable to rapid deployment in response to a pandemic. Overall, our results demonstrate how one could increase the efficacy of single-domain binders toward targets such as SARS-CoV-2 by specifically selecting for binders within an epitope of interest and linking them together to improve potency by avidity. This approach could be applied to target future neutralizing viral epitopes as well as against other protein interfaces of interest.

## Acknowledgements

We thank members of the Wells Lab, particularly those working on COVID-19 projects for their efforts and contributions. Specifically, we would like to thank Dr. Josef Gramespacher for purifying binders and Dr. Matthew Nix (in Dr. Arun Wiita’s Lab) for guidance on pseudotyped virus neutralization assays. Additionally, we thank the entire QCRG for their rapid large-scale collaborative effort. Specifically, we thank Dr. Cristina Puchades and Dr. Caliegh Azumaya for their efforts in optimization of cryoEM grid freezing and data collection; Dr. Tristan Owens for assistance in protein purification; Devan Diwanji for expression of S_ecto_; and Dr. Aashish Manglik for advice on cryoEM experiments and protein purification. We also thank the laboratory of Dr. Peter Kim (Stanford University) for providing plasmids for pseudotyped virus production. Lastly, we thank Dr. Michael Wilson, Dr. Charles Chiu, and Rita Loudermilk of UCSF as well as the patients, for providing convalescent sera.

J.A.W. is supported by generous grants from NCI (R35 GM122451-01), Chan-Zuckerberg Biohub, and UCSF Program for Breakthrough Biomedical Research (PBBR). S.A.L. is a Merck Fellow of the Helen Hay Whitney Foundation. K.S. is a Fellow of the Helen Hay Whitney Foundation. X.X.Z is a Merck Fellow of the Damon Runyon Cancer Research Foundation, DRG-2297-17. J.R.B. and J.Z. are supported by a National Institutes of Health National Cancer Institute F32 5F32CA239417 (to J.R.B,) and 5F32CA236151-02 (to J.Z.). S.Z. is supported by a National Institutes of Health National Cancer Institute T32 (HL007185). N.J.R and I.L are supported by the National Science Foundation (GRFP). D.P.N was the Connie and Bob Lurie Fellow of the Damon Runyon Cancer Research Foundation (DRG-2204-14) and supported by a UCSF-PBBR Postdoctoral Independent Research Award, which is partially funded by the Sandler Foundation.

## QCRG Structural Biology Consortium

In addition to those listed explicitly in the author contributions, the structural biology portion of this work was performed by the QCRG (<u>Q</u>uantitative Biosciences Institute <u>C</u>oronavirus <u>R</u>esearch <u>G</u>roup) Structural Biology Consortium. Listed below are the contributing members of the consortium listed by teams in order of team relevance to the published work. Within each team the team leads are italicized (responsible for organization of each team, and for the experimental design utilized within each team), then the rest of team members are listed alphabetically. **CryoEM grid freezing/collection team:***Caleigh M. Azumaya, Cristina Puchades, Ming Sun*, Julian R. Braxton, Axel F. Brilot, Meghna Gupta, Fei Li, Kyle E. Lopez, Arthur Melo, Gregory E. Merz, Frank Moss, Joana Paulino, Thomas H. Pospiech, Jr., Sergei Pourmal, Alexandrea N. Rizo, Amber M. Smith, Paul V. Thomas, Feng Wang, Zanlin Yu. **CryoEM data processing team:***Miles Sasha Dickinson, Henry C. Nguyen*, Daniel Asarnow, Julian R. Braxton, Melody G. Campbell, Cynthia M. Chio, Un Seng Chio, Devan Diwanji, Bryan Faust, Meghna Gupta, Nick Hoppe, Mingliang Jin, Fei Li, Junrui Li, Yanxin Liu, Gregory E. Merz, Joana Paulino, Thomas H. Pospiech, Jr., Sergei Pourmal, Smriti Sangwan, Raphael Trenker, Donovan Trinidad, Eric Tse, Kaihua Zhang, Fengbo Zhou. **Mammalian cell expression team:***Christian Billesboelle, Melody G. Campbell, Devan Diwanji, Carlos Nowotny, Amber M. Smith, Jianhua Zhao*, Caleigh M. Azumaya, Alisa Bowen, Nick Hoppe, Yen-Li Li, Phuong Nguyen, Cristina Puchades, Mali Safari, Smriti Sangwan, Kaitlin Schaefer, Raphael Trenker, Tsz Kin Martin Tsui, Natalie Whitis. **Protein purification team**: *Daniel Asarnow, Michelle Moritz, Tristan W. Owens, Sergei Pourmal*, Caleigh M. Azumaya, Cynthia M. Chio, Bryan Faust, Meghna Gupta, Kate Kim, Joana Paulino, Jessica K. Peters, Kaitlin Schaefer, Tsz Kin Martin Tsui. **Crystallography team:***Nadia Herrera, Huong T. Kratochvil, Ursula Schulze-Gahmen, Michael C. Thompson, Iris D. Young*, Justin Biel, Ishan Deshpande, Xi Liu. **Bacterial expression team**: *Amy Diallo, Meghna Gupta*, *Erron W. Titus*, Jen Chen, Roberto Efraín Díaz, Loan Doan, Sebastian Flores, Mingliang Jin, Huong T. Kratochvil, Victor L. Lam, Yang Li, Megan Lo, Gregory E. Merz, Joana Paulino, Aye C. Thwin, Zanlin Yu, Fengbo Zhou, Yang Zhang. **Infrastructure team:**David Bulkley, Arceli Joves, Almarie Joves, Liam McKay, Mariano Tabios, Eric Tse. **Leadership team:***Oren S Rosenberg, Kliment A. Verba*, David A. Agard, Yifan Cheng, James S. Fraser, Adam Frost, Natalia Jura, Tanja Kortemme, Nevan J. Krogan, Aashish Manglik, Daniel R. Southworth, Robert M. Stroud. The QCRG Structural Biology Consortium has received support from: Quantitative Biosciences Institute, Defense Advance Research Projects Agency HR0011-19-2-0020 (to D.A.A. and K.A.V.; B. Shoichet PI), FastGrants COVID19 grant (K.A.Verba PI), Laboratory For Genomics Research (O.S. Rosenberg PI) and Laboratory for Genomics Research LGR-ERA (R.M.Stroud PI). RMS is supported by NIH grants AI 50476, GM24485.

## Author Contributions

C.J.B. and D.P.N. designed and constructed the VH-phage library. C.J.B., S.A.L., N.J.R., J.Z., I.L., J. L., K.P. cloned, expressed, and purified the VH binders and/or antigens. C.J.B., S.A.L., J.L. designed and/or conducted the in vitro characterization of the VH binders. J.R.B. designed and conducted the competition ELISA assay with convalescent patient sera. P.S., N.J.R., C.J.B., S.A.L. designed and conducted the pseudovirus neutralization assays. P.S., N.J.R., B.S.Z. designed and conducted the live virus neutralization assays. K.S. prepared samples for cryo-EM and the QCRG SBC performed cryo-EM data collection and analyses. S.A.L., X.X.Z., K.K.L., J.A.W. led coordination of external collaborations and supervised the project. C.J.B., S.A.L., J.A.W. co-wrote the manuscript with input from all the authors.

## Notes

Authors declare that there is no conflict of interest.

## Materials and Methods

### VH Library Construction and Validation by NGS

The VH-phage library was created through bivalent display of VH on the surface of M13 bacteriophage as has been previously described.^30,46,47^ In brief, the DNA phagemid library was created through oligonucleotide mutagenesis. First, the human VH-4D5 sequence was modified with five mutations (35G/39R/45E/47L/50R) in the framework and with restriction sites in each of the CDRs: AgeI in CDR H1, NcoI in CDR H2, and XhoI in CDR H3 (**Table S1**). Oligonucleotides were synthesized by a custom Trimer Phosphoramidite mix for CDR H1 and CDR H2 (Twist Bioscience) and CDR H3 (Trilink Biotechnologies, Inc.) (**Table S2**). After mutagenesis DNA sublibrary pools were digested with appropriate restriction enzymes to remove phagemid template before transformation into SS320 electrocompetent cell (Lucigen) for phage production. NGS of the CDR H3 was performed on the pooled library by amplifying the phagemid from boiled phage with in-house primers. Samples were submitted for analysis on a HiSeq4000 (Illumina) with a custom primer: TGAGGACACTGCCGTCTATTATTGTGCTCGC (Tm = 67 ◦C, GC% = 52). NGS analysis of output was performed using an in-house informatics pipeline written in R. In brief, the raw NGS data sequencing file (*.fastq.gz) was converted into a table comprised of the DNA sequences, the amino acid sequences (CDR H3), and the counts/frequency as columns and then saved as a *.csv file for further analysis (e.g., calculation of: amino acid abundancy, sequence logo, H3 length distribution, etc.). Several filters were applied: i) low-quality sequences containing “N” were removed, ii) sequences with any stop codon were removed; iii) only the sequences that were in-frame were kept. Scripts are available for download at: https://github.com/crystaljie/VH_library_CDR_H3_NGS_analysis_Cole.Bracken.git.

### Cloning, Protein Expression, and Purification

Spike-RBD-monomer, Spike-RBD-Fc, Spike ectodomain (S_ecto_), and ACE2-Fc were produced as biotinylated proteins as previously described.^34^ VH were subcloned from the VH-phagemid into an *E. coli* expression vector pBL347. VH_2_ and VH_3_ were cloned into pBL347 with a 20-amino acid Gly-Ser linker. VH-Fc were cloned into a pFUSE (InvivoGen) vector with a human IgG1 Fc domain. All constructs were sequence verified by Sanger sequencing. VH, VH_2_, and VH_3_ constructs were expressed in *E. coli* C43(DE3) Pro + using an optimized autoinduction media and purified by protein A affinity chromatography similarly to Fabs.^30^ VH-Fc were expressed in Expi293 BirA cells using transient transfection (Expifectamine, Thermo Fisher Scientific). 4 days after transfection, media was harvested, and VH-Fc were purified using protein A affinity chromatography. All proteins were buffer exchanged into PBS by spin concentration and stored in aliquots at −80°C. Purity and integrity of proteins were assessed by SDS-PAGE. All proteins were endotoxin removed using an endotoxin removal kit (Thermo Fischer) prior to use in neutralization assays.

### Phage selection with VH-phage library

Phage selections were done according to previously established protocols.^30^ Selections were performed using biotinylated antigens captured with streptavidin-coated magnetic beads (Promega). In each round, the phage pool was first cleared by incubation with beads loaded with 500 nM ACE2-Fc/Spike-RBD-Fc complex. The unbound phage were then incubated with beads loaded with Spike-RBD-Fc. After washing, the bound phage was eluted by the addition of 2 μg/mL of TEV protease. In total, four rounds of selection were performed with decreasing amounts of Spike-RBD-Fc as indicated in Figure 2A. All steps were done in PBS buffer + 0.02% Tween-20 + 0.2% BSA (PBSTB). Individual phage clones from the third and fourth round of selections were analyzed by phage ELISA.

### Phage ELISA

For each phage clone, 4 different conditions were tested—Direct: Spike-RBD-Fc, Competition: Spike-RBD-Fc with equal concentration of Spike-RBD-Fc in solution, Negative selection: ACE2-Fc/Spike-RBD-Fc complex, and Control: Fc. 384-well Nunc Maxisorp flat-bottom clear plates (Thermo Fisher Scientific) were coated with 0.5 μg/mL of NeutrAvidin in PBS overnight at 4°C and subsequently blocked with PBS + 0.02% Tween-20 + 2% BSA for 1 hr at room temperature. Plates were washed 3X with PBS containing 0.05% Tween-20 (PBST) and were washed similarly between each of the steps. 20 nM of biotinylated Spike-RBD-Fc, ACE2-Fc/Spike-RBD-Fc complex, or Fc diluted in PBSTB was captured on the NeutrAvidin-coated wells for 30 min, then blocked with PBSTB + 10 μM biotin for 30 min. Phage supernatant diluted 1:5 in PBSTB were added for 20 min. For the competition samples, the phage supernatant was diluted into PBSTB with 20 nM Spike-RBD-Fc. Bound phage were detected by incubation with anti-M13-HRP conjugate (Sino Biological)(1:5000) for 30 min, followed by addition of TMB substrate (VWR International). The reaction was quenched with addition of 1 M phosphoric acid and the absorbance at 450 nm was measured using a Tecan M200 Pro spectrophotometer.

### ELISA EC50 with Spike-RBD-monomer

384-well Nunc Maxisorp flat-bottom clear plates were prepared similarly to the phage ELISA protocol (above) by coating with neutravidin, followed by blocking with PBST + 2% BSA, incubation with 20 nM Spike-RBD-monomer, and blocking by PBSTB + 10 μM biotin. VH binders in 4-fold dilutions ranging from 500 nM to 2.8 pM were added for 1 hour. Bound VH was detected by incubation with Protein A HRP conjugate (Thermo Fischer Scientific) (1:10,000) for 30 min, followed by addition of TMB substrate for 5 min, quenching by 1 M phosphoric acid, and detection of absorbance at 450 nm. Each concentration was tested in duplicate, and the assay was repeated three times.

### Bio-layer interferometry (BLI) Experiments

Bio-layer interferometry data (BLI) were measured using an Octet RED384 (ForteBio) instrument. Spike-RBD or S_ecto_ were immobilized on a streptavidin or Ni-NTA biosensor and loaded until 0.4 nm signal was achieved. After blocking with 10 μM biotin, purified binders in solution was used as the analyte. PBSTB was used for all buffers. Data were analyzed using the ForteBio Octet analysis software and kinetic parameters were determined using a 1:1 monovalent binding model.

### Competition ELISA with COVID-19 convalescent patient sera

Competition ELISA with convalescent patient sera was conducted with the same patient sera as previously reported.^36^ Samples were collected in accordance with the Declaration of Helsinki using protocols approved by the UCSF Institutional Review Board (Protocol 20-30338). Patients were voluntarily recruited based on their history of prior SARS-CoV-2 infection. All patients provided written consent. Patient sera were de-identified prior to delivery to the Wells Lab, where all experiments presented here were performed. Briefly, sera were obtained as described from patients with a history of positive nasopharyngeal SARS-CoV-2 RT-PCR test and at least 14 days after the resolution of their COVID-19 symptoms. Healthy control serum was obtained prior to the emergence of SARS-CoV-2. Sera were heat-inactivated (56°C for 60 min) prior to use. Competitive serology using biotinylated SARS-CoV-2 Spike-RBD as the capture antigen was performed as previously reported with slight modifications.^36^ Instead of supplementing sera diluted 1:50 in 1% nonfat milk with 100 nM ACE2-Fc, 100 nM of each of the indicated VH-Fc fusions was used. Bound patient antibodies were then detected using Protein L-HRP (Thermo Fisher Scientific 32420, [1:5000]). Background from the raw ELISA signal in serum-treated wells was removed by first subtracting the signal measured in NeutrAvidin-alone coated wells then subtracting the signal detected in antigen-coated wells incubated with 1% nonfat milk + 100 nM competitor.

### Differential Scanning Fluorimetry (DSF)

DSF was conducted as previously described.^30^ Briefly, purified protein was diluted to 0.5 μM or 0.25 μM in buffer containing Sypro Orange 4x (Invitrogen) and PBS and assayed in a 384-well white PCR plate. All samples were tested in duplicate. In a Roche LC480 LightCycler, the sample was heated from 30°C to 95°C with a ramp rate of 0.3°C per 30 sec and fluorescence signal at 490 nm and 575 nm were continuously collected. T_m_ was calculated using the Roche LC480 LightCycler software.

### Size Exclusion Chromatography (SEC)

SEC analysis was performed using an Äkta Pure system (GE Healthcare) using a Superdex 200 Increase 10/300 GL column. 100 μL of 2-3 mg/mL of each analyte was injected and run with a constant mobile phase of degassed 10 mM Tris pH 8.0 200 mM NaCl. Absorbance at 280 nm was measured. The post-lyophilization and reconstitution SEC was performed using an Agilent HPLC 1260 Infinity II LC System using an AdvanceBio SEC column (300 Å, 2.7 μm, Agilent). Fluorescence (excitation 285 nm, emission 340 nm) was measured.

### Preparation of SARS-CoV-2 pseudotyped virus and HEK-ACE2 overexpression cell line

HEK293T-ACE2 cells were a gift from Arun Wiita’s laboratory at the University of California, San Francisco. Cells are cultured in D10 media (DMEM + 1% Pen/Strep + 10% heat-inactivated FBS).

Plasmids to generate pseudotyped virus were a gift from Peter Kim’s lab at Stanford University and SARS-Cov-2 pseudotyped virus was prepared as previously described.^48^ Briefly, plasmids at the designated concentrations were added to OptiMEM media with FuGENE HD Transfection Reagent (Promega) at a 3:1 FuGENE:DNA ratio, incubated for 30 min, and subsequently transfected into HEK-293T cells. After 24 hrs, the supernatant was removed and replaced with D10 culture media. Virus was propagated for an additional 48 hrs, and the supernatant was harvested and filtered. Virus was stored at 4°C for up to 10 days.

HEK-ACE2 were seeded at 10,000 cells/well on 96-well white plates (Corning, cat. 354620). After 24 hrs, pseudotyped virus stocks were titered via a two-fold dilution series in D10 media and 40 μL were added to cells. After 60 hrs, infection and intracellular luciferase signal was determined using Bright-Glo™ Luciferase Assay (Promega), and the dilution achieving maximal luminescent signal within the linear range, ~3-5 x 10^5^ luminescence units, was chosen as the working concentration for neutralization assays.

### Pseudotyped viral neutralization assays

HEK-ACE2 were seeded at 10,000 cells/well in 40 μL of D10 on 96-well white plates (Corning, cat. 354620) 24 hours prior to infection. To determine IC_50_ for pseudotyped virus, dose series of each VH binder were prepared at 3x concentration in D10 media and 50 μL were aliquoted into each well in 96-well plate format. Next, 50 μL of virus were added to each well, except no virus control wells, and the virus and blocker solution was allowed to incubate for 1 hr at 37°C. Subsequent to pre-incubation, 80 μL of the virus and blocker inoculum were transferred to HEK-ACE2. After 60 hrs of infection at 37°C, intracellular luciferase signal was measured using the Bright-Glo™ Luciferase Assay.

### Live Virus Neutralization Assays

Live virus neutralization assays were done as previously described.^34^ All handling and experiments using live SARS-CoV-2 virus clinical isolate 2019-nCoV/USA-WA1/2020 (BEI Resources) was conducted under Biosafety Level 3 containment with approved BUA and protocols. Briefly, VeroE6 cells were cultured in Minimal Essential Media (MEM), 10% FBS, 1% Pen-Strep. Virus was incubated in infection media (EMEM 0% FBS) containing different concentrations of binders for 1 hr at 37°C and subsequently added to VeroE6 cells for a low volume infection for 1 hr. EMEM with 10% FBS was added and cells incubated at 37°C for 16 hrs before RNA harvest. Viral entry into cells and cellular transcription of viral genes was measured by qPCR using the N gene, E gene, RdRp gene and host GUSB and host ACTB as controls. Relative copy number (RCN) of viral transcript level compared to host transcript was determine using the ΔΔCT method.

### Expression and purification of Spike ectodomain for cryo-EM

To obtain pre-fusion spike ectodomain, methods similar to the previous reports were used.^33,49^ The expression plasmid, provided by the McLellan lab, was used in a transient transfection with 100 mL, high-density Chinese Hamster Ovary (ExpiCHO, Thermo Fisher) culture following the “High Titer” protocol provided by Thermo Fisher. Six to nine days post-transfection, the supernatant was collected with centrifugation at 4,000xg at room temperature. The clarified supernatant was then incubated with Ni-Sepharose Excel resin (Cytiva Life Sciences) for ninety minutes at room temperature. After incubation, the nickel resin was washed with 20 mM Tris (pH 8), 200 mM NaCl, and 20 mM imidazole with ten column volumes. Protein was eluted from the nickel resin with 20 mM Tris (pH 8), 200 mM NaCl, and 500 mM imidazole. Eluate was then concentrated with a 50 MWCO Amicon Ultra-15 centrifugal unit by centrifugation at 2500xg, room temperature. The eluate was concentrated, filtered with a 0.2 μm filter, and injected onto a Superose6 10/300 GL column equilibrated with 10 mM Tris (pH 8), 200 mM NaCl. The fractions corresponding to monodisperse spike were collected and the concentration was determined using a nanodrop.

### Cryo-EM sample preparation, data collection and processing

2 μM Spike ectodomain was mixed with 5-fold excess VH_3_ B01 and applied (3 *μ*L) to holey carbon Au 200 mesh 1.2/1.3 Quantifoil grids. Grids were blotted and plunge frozen using Mark 4 Vitrobot (ThermoFisher) at 4°C and 100% humidity, utilizing blot force 0 and blot time of 4 sec. 1656 images were collected on Titan Krios (ThermoFisher) equipped with K3 direct detector operated in CDS mode (Gatan Ametek) and an energy filter (Gatan Ametek) at nominal magnification of 105000x (0.834 Å/physical pixel). Dose fractionated movies were collected with a total exposure of 6 seconds and 0.04 seconds per frame at a dose rate of 9 electrons per physical pixel per second. Movies were corrected for motion and filtered to account for electron damage utilizing MotionCor2.^50^ Drift corrected sums were imported into cryoSPARC2 processing package.^51^ Micrographs were manually curated, CTF was estimated utilizing patches and particles were picked with a Gaussian blob. Previous Spike ectodomain structure was imported as an initial model (low pass filtered to 30 Å) and multiple rounds of 3D and 2D classification were performed. Images were re-picked with the best looking 2D class averages low pass filtered to 30 Å, and multiple rounds of 3D classification were performed again to obtain a homogeneous stack of Spike trimer particles. Majority of the particles went into classes putatively representing excess unbound VH_3_ B01 and the final Spike like particle stack only contained ~21000 particles. Non-uniform homogeneous refinement of the particle stack resulted in global resolution of 3.2 Å (masked) utilizing 0.143 FSC cut off.^52^ PDB:6X2B was rigid body fit into the resulting reconstruction in UCSF Chimera.^53^ The RBDs of two Spike trimers were moved as a rigid body to accommodate the cryoEM density. The cryo-EM reconstruction was low pass filtered to 6Å to better visualize the VH densities. Homology model was built based on the PDB:4G80 for the VH domains and the resulting and individual VH domains were rigid body fit into the 6Å cryoEM density as depicted on **Fig. 5A**. The resulting model was relaxed into the cryoEM map low pass filtered to 6Å with Rosetta FastRelax protocol. For **Fig. S13** the local resolution was estimated using ResMap.^54^ The final figures were prepared using ChimeraX.^55^

