## Supplementary Material for "Bi-paratopic and multivalent human VH domains neutralize SARS-CoV-2 by targeting distinct epitopes within the ACE2 binding interface of Spike"

**Authors and Affiliations:**

Colton J. Bracken[1], Shion A. Lim[1], Paige Solomon[1], Nicholas J. Rettko[1], Duy P. Nguyen[1], Beth Shoshana Zha[2], Kaitlin Schaefer[1], James R. Byrnes[1], Jie Zhou[1], Irene Lui[1], Jia Liu[1], Katarina Pance[1], QCRG Structural Biology Consortium[3,#], Xin X. Zhou[1], Kevin K. Leung[1], James A. Wells[1,4,*]

**Table S1: Modified VH-4D5 template sequence used for phagemid construction**

Green codons represent amino acid changes (35G/39R/45E/47L/50R) in the framework. Restriction sites in each of the CDRs: AgeI in CDR H1 (blue), NcoI in CDR H2 (purple), and XhoI in CDR H3 (red).

| | |
|---|---|
| VH-4D5 template | 5’-gaggttcagctggtggagtctggcggtggcctggtgcagccagggggctcactccgtttgtcctgtgcagcttctg gcttcAACATCaccggtACTTATATCggctgggtgcgtcgcgccccgggtaagggcgaggaactggttgcacgtatc TACCCCACGccatggTATACCCGCtatgccgatagcgtcaagggccgtttcactataagcgcagacacatccaaa aacacagcctacctacaaatgaacagcttaagagctgaggacactgccgtctattattgtgctcgcTGGGGAGGGGA CGGATTCTACctcgaggactactggggtcaaggaaccctggtcaccgtctcctcg-3’ |

**Table S2: Oligonucleotides used for VH library construction denoted by their length**

XXX and ZZZ are representative codons that correspond to different amino acid frequencies previously specified in **Fig. 1B**. S = C/G, W = A/T, K = G/T.

| | |
|---|---|
| H1-5 | 5'–cctgtgcagcttctggcttc ZZZ ZZZ ZZZ ZZZ ZZZ ZZZ atcggctgggtgcgtcg-3' |
| H1-6 | 5'–cctgtgcagcttctggcttc ZZZ ZZZ ZZZ ZZZ ZZZ ZZZ ZZZ atcggctgggtgcgtcg-3' |
| H1-7 | 5'–cctgtgcagcttctggcttc ZZZ ZZZ ZZZ ZZZ ZZZ ZZZ ZZZ ZZZ atcggctgggtgcgtcg-3' |
| H2 | 5'–gcgaggaactggttgcacgtatc ZZZ ZZZ ZZZ ZZZ ZZZ ZZZ ZZZ ZZZ tatgccgatagcgtcaagggcc-3' |
| H3-4 | 5'– c gtc tat tat tgt gct cgc XXX XXX gSt WtK gac tac tgg ggt caa gg -3' |
| H3-5 | 5'– c gtc tat tat tgt gct cgc XXX XXX XXX gSt WtK gac tac tgg ggt caa gg -3' |
| H3-6 | 5'– c gtc tat tat tgt gct cgc XXX XXX XXX XXX gSt WtK gac tac tgg ggt caa gg -3' |
| H3-7 | 5'– c gtc tat tat tgt gct cgc XXX XXX XXX XXX XXX gSt WtK gac tac tgg ggt caa gg -3' |
| H3-8 | 5'– c gtc tat tat tgt gct cgc XXX XXX XXX XXX XXX XXX gSt WtK gac tac tgg ggt caa gg -3' |
| H3-9 | 5'– c gtc tat tat tgt gct cgc XXX XXX XXX XXX XXX XXX XXX gSt WtK gac tac tgg ggt caa gg -3' |
| H3-10 | 5'– c gtc tat tat tgt gct cgc XXX XXX XXX XXX XXX XXX XXX XXX gSt WtK gac tac tgg ggt caa gg -3' |
| H3-11 | 5'– c gtc tat tat tgt gct cgc XXX XXX XXX XXX XXX XXX XXX XXX XXX gSt WtK gac tac tgg ggt caa gg -3' |
| H3-12 | 5'– c gtc tat tat tgt gct cgc XXX XXX XXX XXX XXX XXX XXX XXX XXX XXX gSt WtK gac tac tgg ggt caa gg -3' |
| H3-13 | 5'– c gtc tat tat tgt gct cgc XXX XXX XXX XXX XXX XXX XXX XXX XXX XXX XXX gSt WtK gac tac tgg ggt caa gg -3' |
| H3-14 | 5'– c gtc tat tat tgt gct cgc XXX XXX XXX XXX XXX XXX XXX XXX XXX XXX XXX XXX gSt WtK gac tac tgg ggt caa gg -3' |
| H3-15 | 5'– c gtc tat tat tgt gct cgc XXX XXX XXX XXX XXX XXX XXX XXX XXX XXX XXX XXX XXX gSt WtK gac tac tgg ggt caa gg -3' |
| H3-16 | 5'– c gtc tat tat tgt gct cgc XXX XXX XXX XXX XXX XXX XXX XXX XXX XXX XXX XXX XXX XXX gSt WtK gac tac tgg ggt caa gg -3' |
| H3-17 | 5'– c gtc tat tat tgt gct cgc XXX XXX XXX XXX XXX XXX XXX XXX XXX XXX XXX XXX XXX XXX XXX gSt WtK gac tac tgg ggt caa gg -3' |
| H3-18 | 5'– c gtc tat tat tgt gct cgc XXX XXX XXX XXX XXX XXX XXX XXX XXX XXX XXX XXX XXX XXX XXX XXX gSt WtK gac tac tgg ggt caa gg -3' |

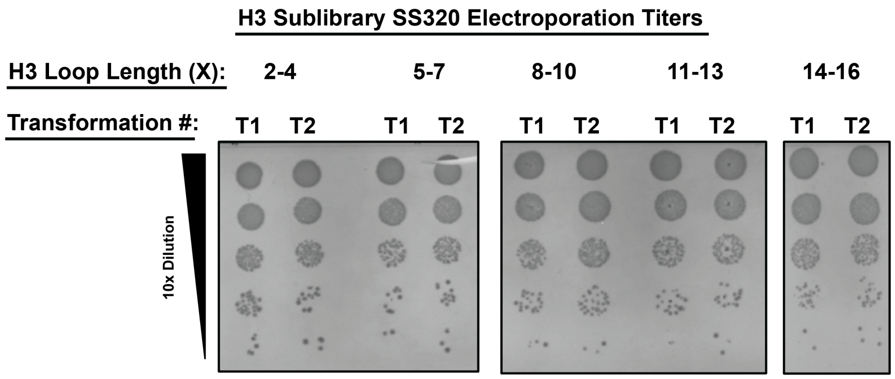


| Sublibrary | Transformants (cfu) |
|---|---|
| 2-4 | $1.33 \times 10^{10}$ |
| 5-7 | $8.33 \times 10^{9}$ |
| 8-10 | $1.16 \times 10^{10}$ |
| 11-13 | $4.99 \times 10^{9}$ |
| 14-16 | $1.16 \times 10^{10}$ |
| Total: | $4.98 \times 10^{10}$ |

**Figure S1: VH-Phage sublibrary construction**

Titers from electroporated SS320 competent cells for each H3 length sublibrary are shown. Final number of transformants (cfu) was calculated as the sum from each sublibrary.

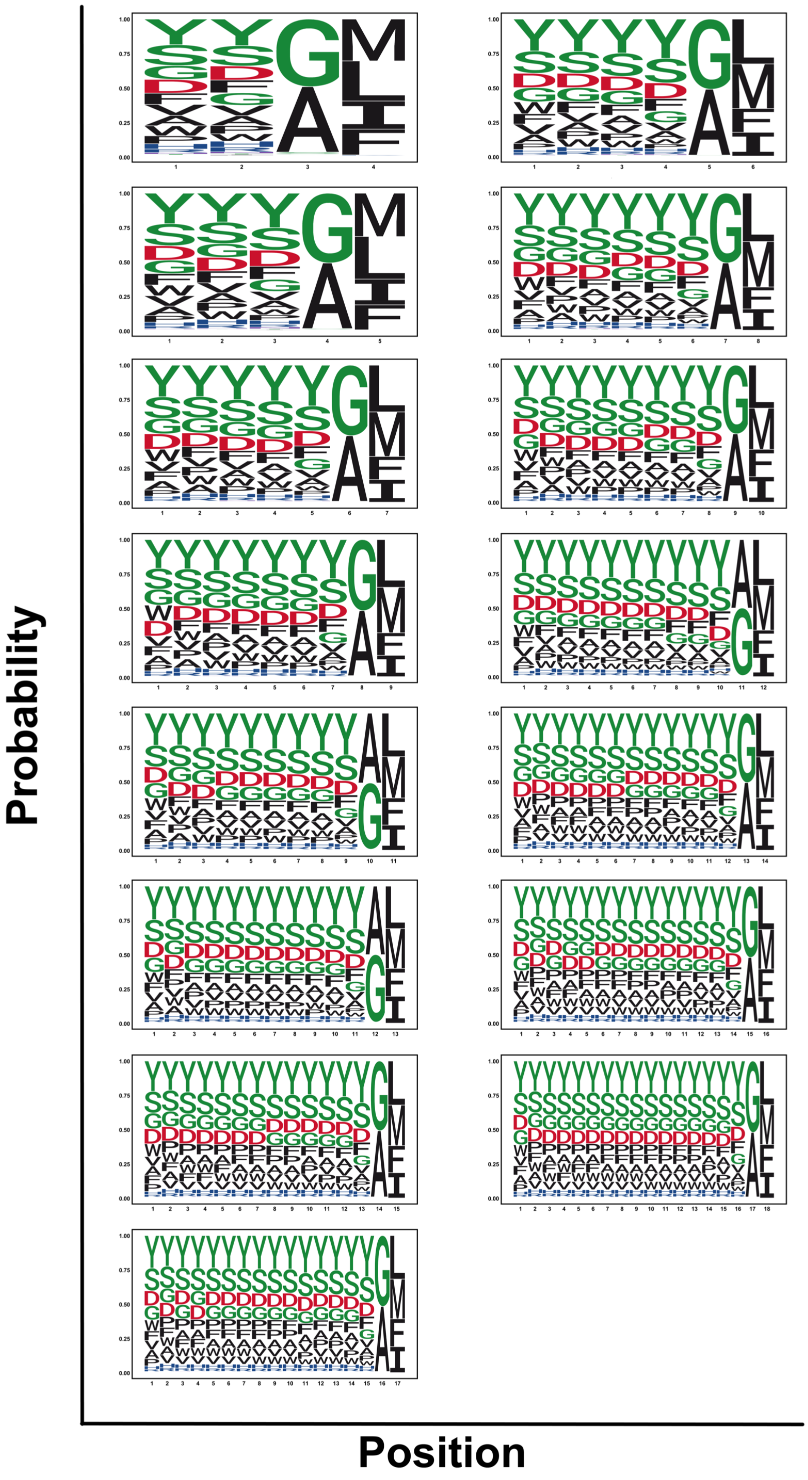


**Figure S2: Positional amino acid analysis of all CDR H3 loops**

NGS analysis shows amino acid composition of the H3 loops (X = 1-18) of unique clones. Plots show positional frequency distribution matches designed frequencies. Position 1 refers to residue 95 (Kabat definition).

### Round 3 Enrichment

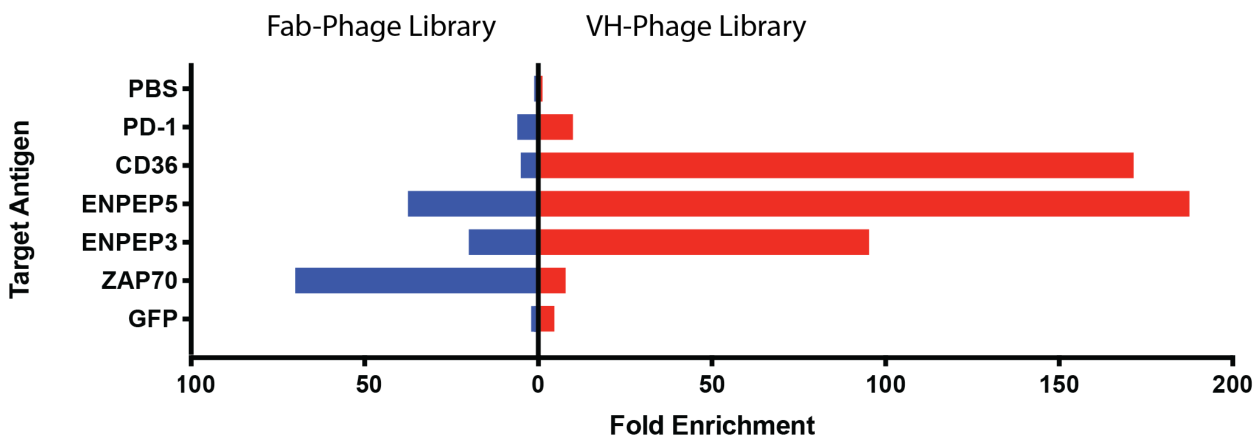


**Figure S3: Enrichment comparison between Fab-phage library and VH-phage library**

Phage-selections were performed at the same time on a representative group of antigens. Antigens included both cytosolic and membrane proteins. Individual fold enrichment values were calculated by comparing phage titers at round three to, Fc-biotin, which was used to clear the phage pool prior to each round of selection. Phosphate buffered saline (PBS) was included as a control.

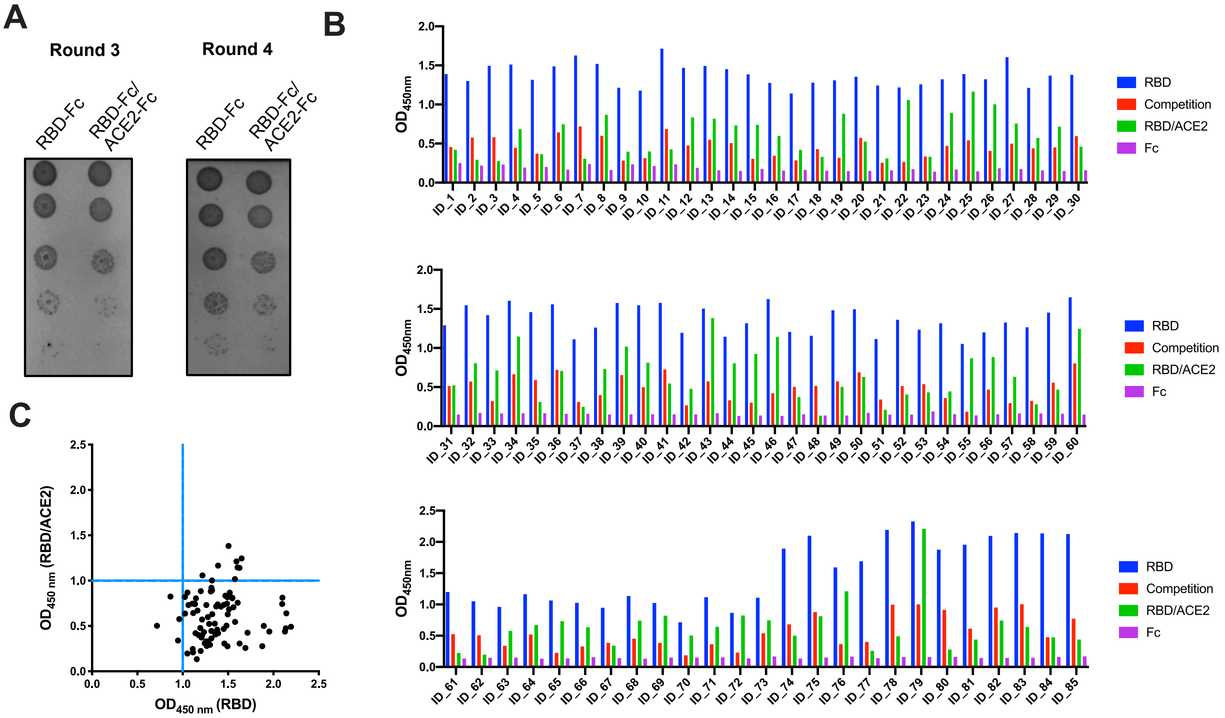


**Figure S4: Phage selection with VH-phage library and characterization of unique clones**

**(A)** Titer of VH-phage after 3 or 4 rounds of selection show significant enrichment for VH-phage that bind Spike-RBD-Fc over VH-phage that bind Spike-RBD-Fc/ACE2-Fc complex. (**B**) ELISA of unique VH-phage against Spike-RBD-Fc, competition with soluble Spike-RBD-Fc, Spike-RBD-Fc/ACE2-Fc complex, or Fc antigen. (**C**) Plot of ELISA data for Spike-RBD-Fc alone vs Spike-RBD-Fc/ACE2-Fc complex shows that most of the unique VH-phage recognize the unmasked Spike-RBD-Fc antigen, suggesting they bind the same epitope as ACE2.

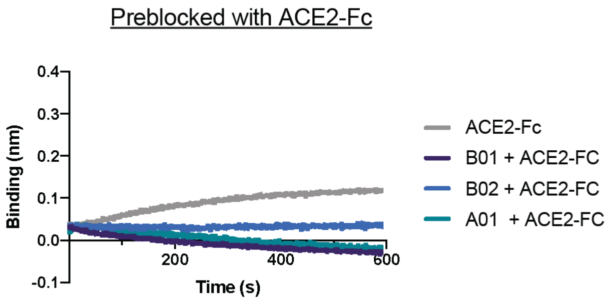


**Figure S5: Epitope binning of lead VH domain binders with ACE2-Fc on Spike-RBD**

Sequential bio-layer interferometry (BLI) binding traces. Spike-RBD was loaded onto sensor tip and preblocked with ACE2-Fc, followed by adding either VH A01, VH B01, or VH B02. These second association data indicate that ACE2-Fc prevents binding of either of the three VH domains, suggesting they share the same binding epitope on Spike-RBD.

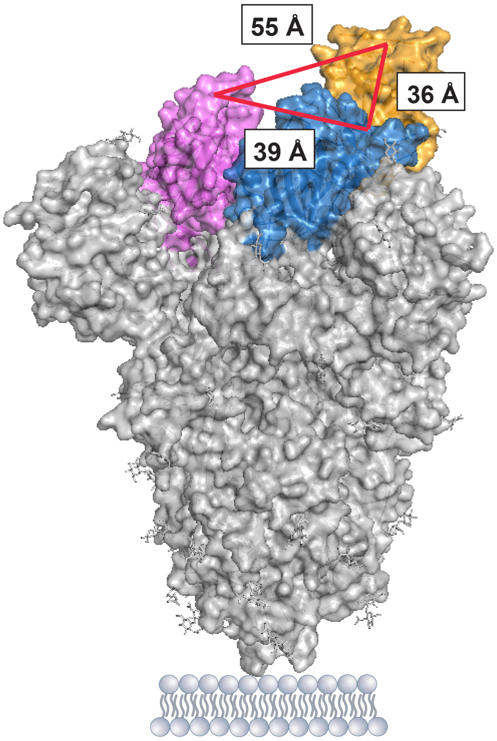


**Figure S6: Measurements of inter-RBD distance on Spike trimer**

Structure of SARS-CoV-2 Spike trimer (PDB: 6VSB)[1]. The three RBDs are shown in color. RBD colored yellow is in the “up” position while RBDs colored in blue and pink are in the “down” position. Distance between the mid-points of the ACE2 binding interface (PDB: 6M17)[2] on respective RBDs was measured in Pymol.

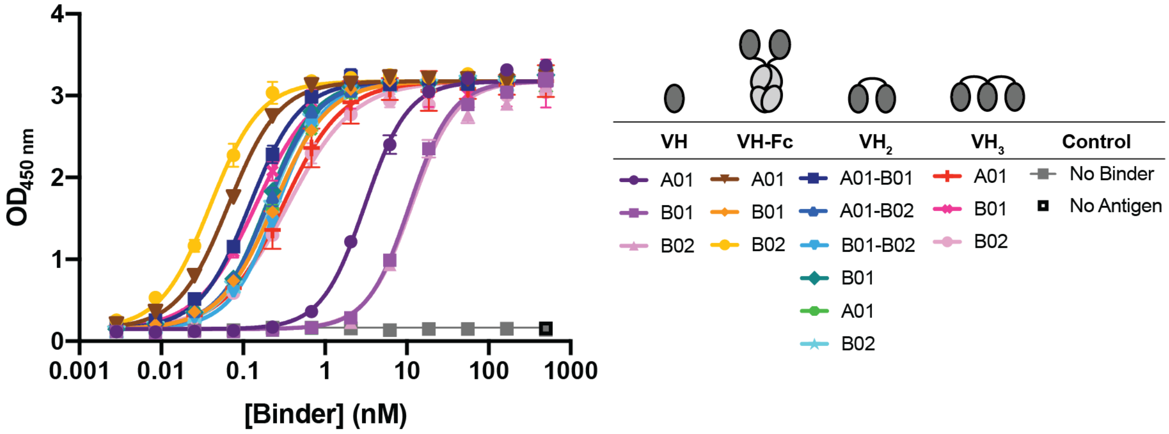


**Figure S7: ELISA of VH binders to Spike-RBD**

ELISA of VH binders against Spike-RBD. Data presented show the average and standard deviation from three independent experiments. Data were fit to a non-linear, four-parameter variable slope regression model using Prism 8 to obtain EC50 values for each binder (**Table S3**).

**Table S3: ELISA EC50 of VH Binders**

| VH ID | EC50 (nM) |
|---|---|
| VH A01 | 3.55 ± 0.67 |
| VH B01 | 12.0 ± 1.6 |
| VH B02 | 14.5 ± 4.3 |
| VH-Fc A01 | 0.0738 ± 0.011 |
| VH-Fc B01 | 0.265 ± 0.042 |
| VH-Fc B02 | 0.0415 ± 0.002 |
| $VH^2$ A01-B01 | 0.139 ± 0.027 |
| $VH^2$ A01-B02 | 0.208 ± 0.014 |
| $VH^2$ B01-B02 | 0.295 ± 0.049 |
| $VH^2$ A01 | 0.280 ± 0.067 |
| $VH^2$ B01 | 0.230 ± 0.053 |
| $VH^2$ B02 | 0.217 ± 0.001 |
| $VH^3$ A01 | 0.323 ± 0.023 |
| $VH^3$ B01 | 0.135 ± 0.011 |
| $VH^3$ B02 | 0.328 ± 0.032 |

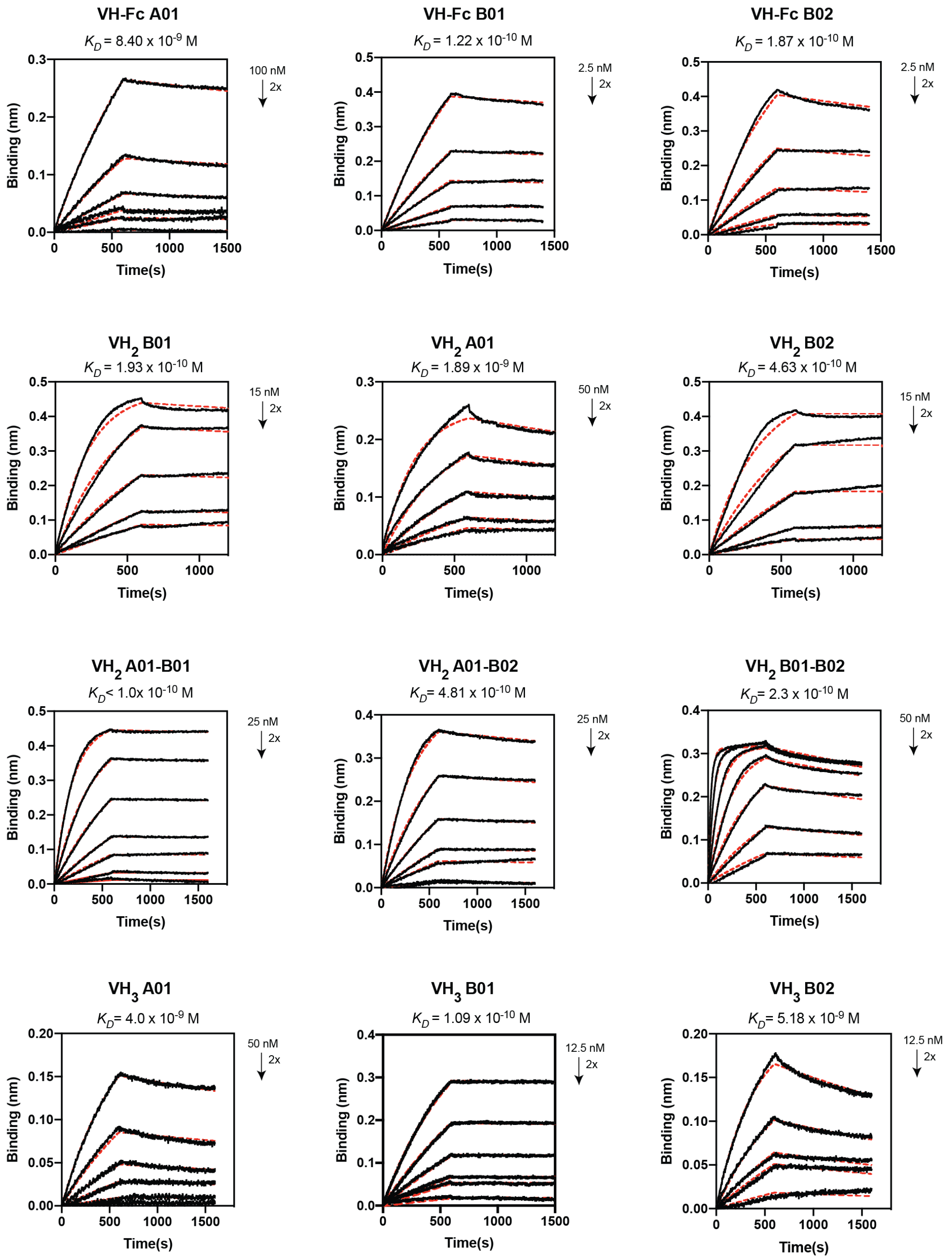


**Figure S8: Multipoint Biolayer Interferometry (BLI) of VH Binders to Spike-RBD**

Multipoint BLI experiments for each VH binder. The data were fit to a 1:1 binding model using the Octet ForteBio software to determine the binding affinity for the measured interaction.

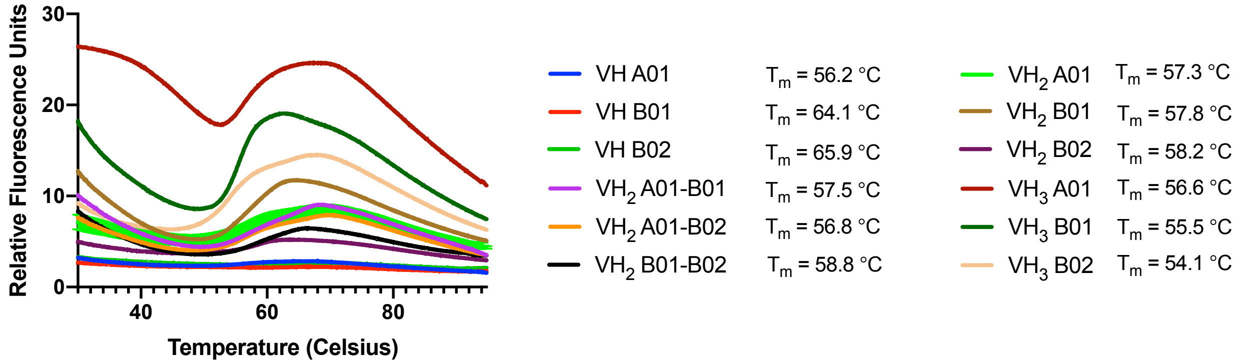


**Figure S9: Differential Scanning Fluorimetry (DSF) of VH Binders**

The melting temperature ($T_m$) of each VH binder was determined by differential scanning fluorimetry. Data and $T_m$ presented are an average of two replicates.

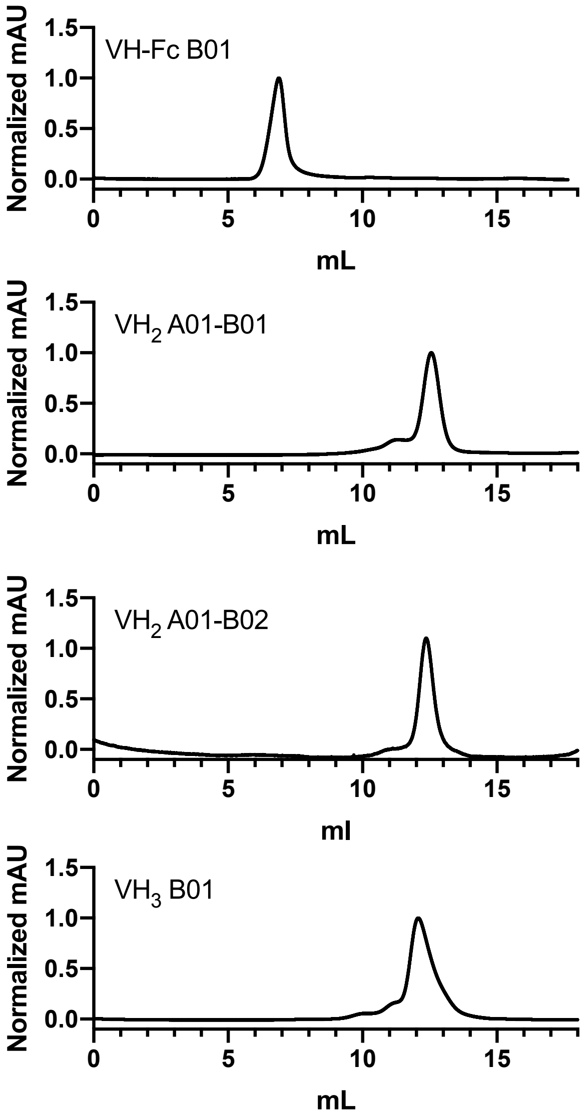


**Figure S10: Size Exclusion Chromatography of VH Binders**

Size exclusion chromatography (SEC) traces of VH-Fc B01, $VH_2$ A01-B01, $VH_2$ A01-B02, and $VH_3$ B01 on a Superdex 200 Increase 10/300 GL column.

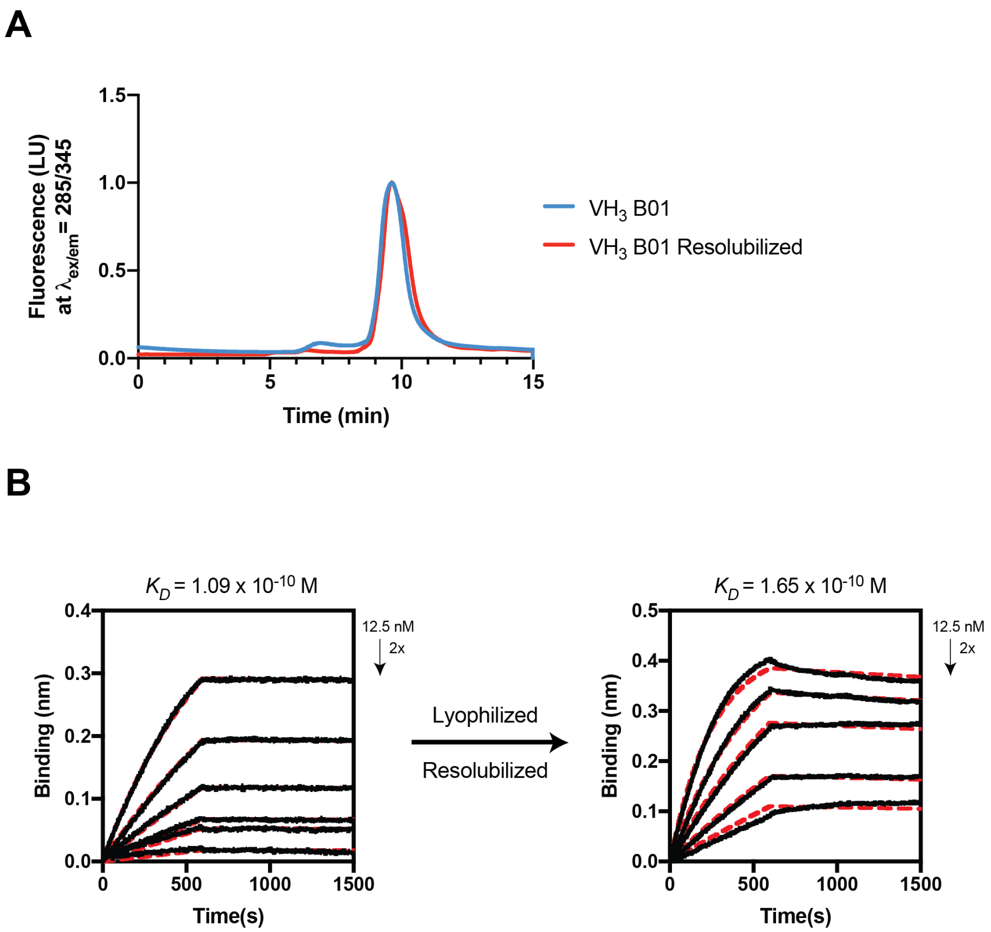


**Figure S11: Lyophilization and reconstitution of $VH_3$ B01**

**(A)** SEC trace of $VH_3$ B01 on an AdvanceBio 300 Å column before and after lyophilization shows that the elution profile of $VH_3$ B01 is unchanged. (**B**) Multipoint BLI of $VH_3$ B01 binding to Spike-RBD before and after lyophilization and resolubilization show that the binding affinity of $VH_3$ B01 is largely unchanged.

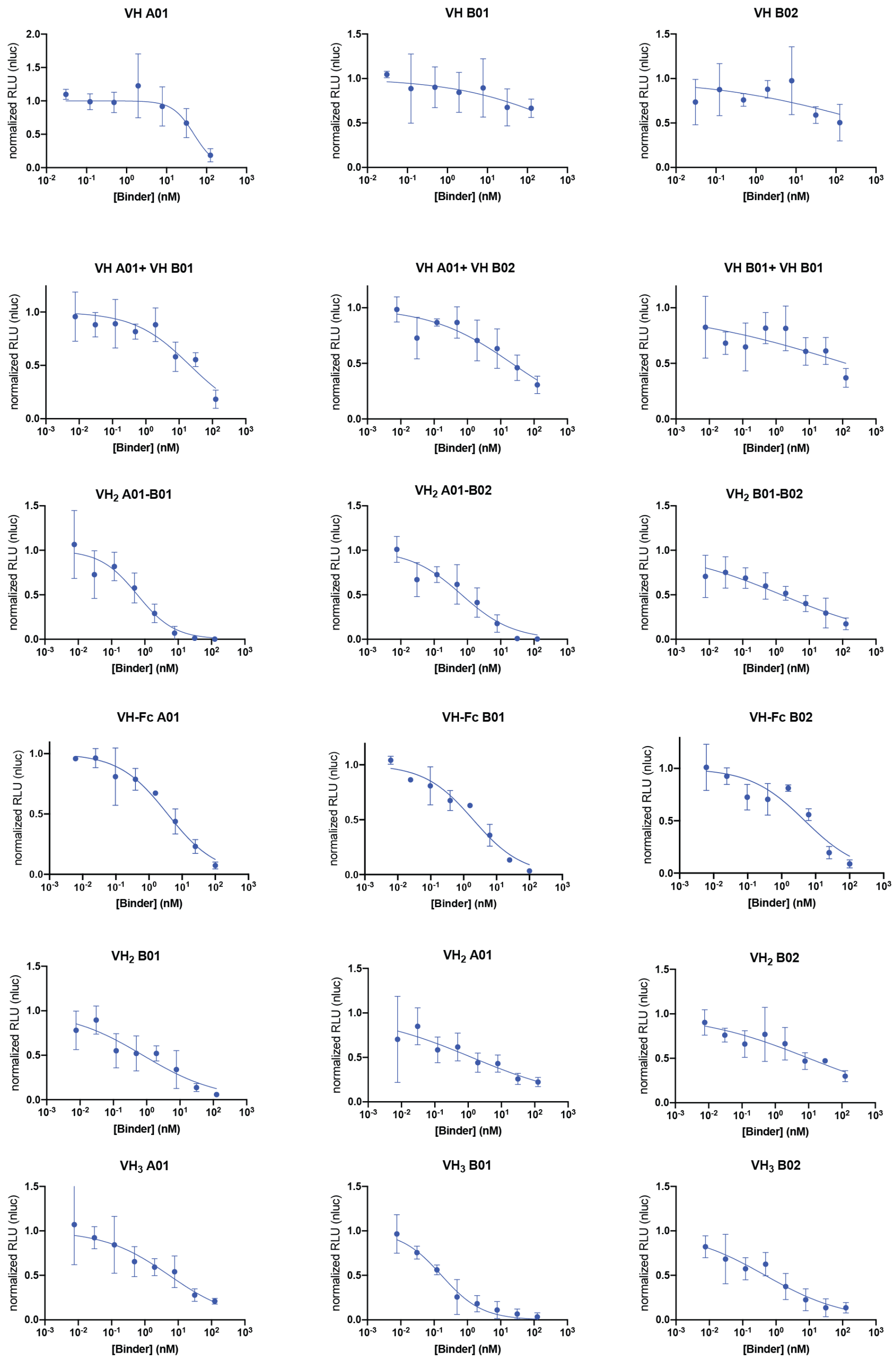


**Figure S12: Pseudotyped virus neutralization by VH Binders**

Pseudotyped virus neutralization assays of VH binders. Data represent average and standard deviation of two or three biological replicates. Data were fit to a non-linear, four-parameter variable slope regression model using Prism 8 to obtain $IC_{50}$ values.

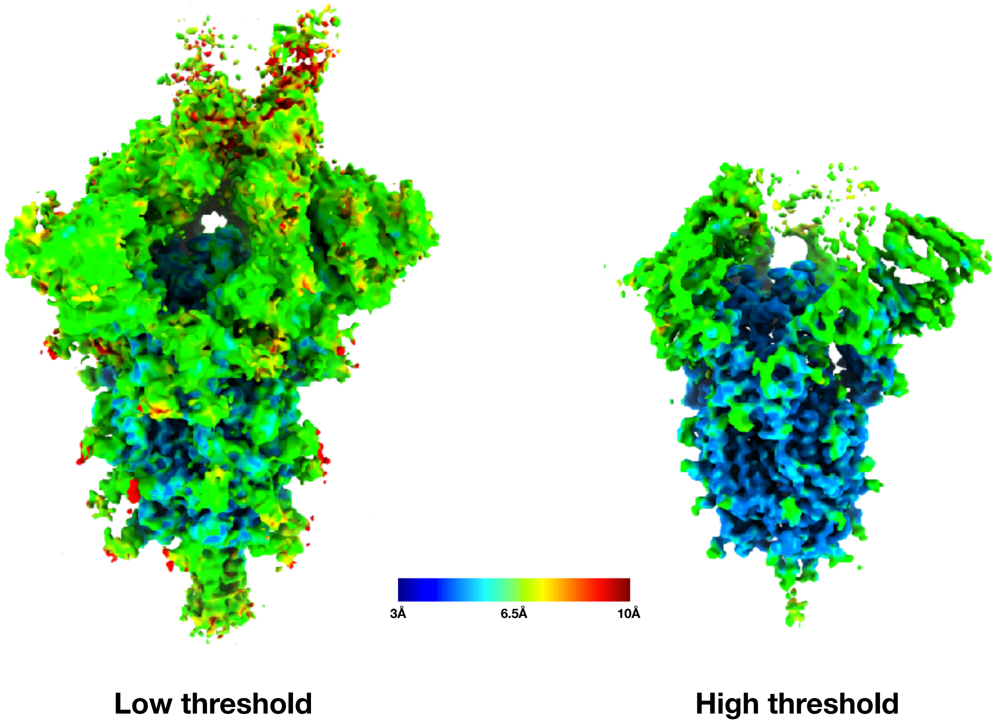


**Figure S13: Cryo-EM reconstruction of SARS-CoV-2 Spike trimer**

The SARS-CoV-2 Spike trimer + $VH_3$ B01 cryo-EM reconstruction from non-uniform refinement in cryoSPARC at two different thresholds colored by resolution in the range from 3 Å to 10 Å. At high threshold the core S2 clearly displays high resolution features but the periphery of the molecule is closer to 6-7 Å.

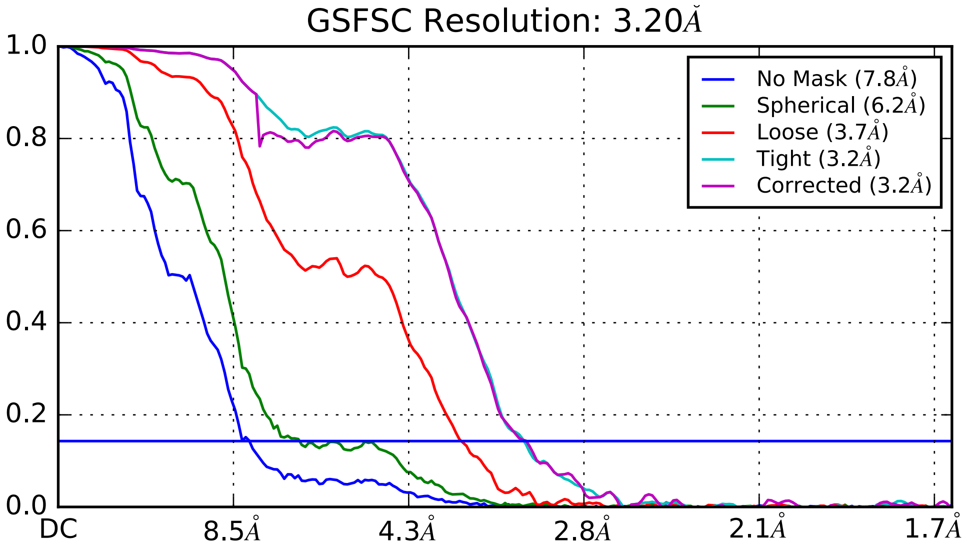


**Figure S14: Gold standard FSC curves from final iteration of non-uniform refinement in cryoSPARC for masked and unmasked reconstructions.**
